# Single-molecule live-cell imaging visualizes parallel pathways of prokaryotic nucleotide excision repair

**DOI:** 10.1101/515502

**Authors:** Harshad Ghodke, Han Ngoc Ho, Antoine M van Oijen

**Affiliations:** Molecular Horizons and School of Chemistry and Molecular Bioscience, University of Wollongong, Wollongong, New South Wales, 2522, Australia; Illawarra Health and Medical Research Institute, Wollongong, New South Wales, 2522, Australia

## Abstract

In the model organism *Escherichia coli*, helix distorting lesions are recognized by the UvrAB damage surveillance complex in the global genomic nucleotide excision repair pathway (GGR). Alternately, during transcription-coupled repair (TCR), UvrA is recruited to Mfd at sites of RNA polymerases stalled or paused by lesions. Ultimately, damage recognition is mediated by UvrA, culminating in the loading of the damage verification enzyme UvrB. We set out to characterize the differences in the kinetics of interactions of UvrA with Mfd and UvrB. We followed functional, fluorescently tagged UvrA molecules in live cells and measured their residence times in TCR-deficient or wild-type cells. We demonstrate that the lifetimes of UvrA in Mfd-dependent or Mfd-independent interactions in the absence of exogenous DNA damage are comparable in live cells, and are governed by UvrB. Upon UV irradiation, we found that the lifetimes of UvrA strongly depended on, and matched those of Mfd. Here, we illustrate a non-perturbative, imaging-based approach to quantify the kinetic signatures of damage recognition enzymes participating in multiple pathways in cells.

## Introduction

Across the various domains of life, the recognition and repair of bulky, helix-distorting lesions in chromosomal DNA is coordinated by nucleotide excision repair (NER) factors. Damage detection occurs in two stages: a dedicated set of damage surveillance enzymes (reviewed in ref.^1, 2, 3^) (namely the prokaryotic UvrA, and the eukaryotic UV-DDB, XPC, XPA and homologs) constantly survey genomic DNA for lesions. At sites of putative DNA damage, these enzymes load specific factors (UvrB in prokaryotes, TFIIH and homologs in eukaryotes) that unwind the DNA and verify the location of the damage with nucleotide resolution (Fig. 1a) (reviewed in ref.^2, 3^). Subsequently, specialized endonucleases (prokaryotic UvrC and homologs, and the eukaryotic XPF-ERCC1/XPG and homologs) are recruited to the site of the DNA, resulting in cleavage of the single-stranded DNA (ssDNA) patch containing the lesion (reviewed in ref.^2, 3^).

**Figure 1:**
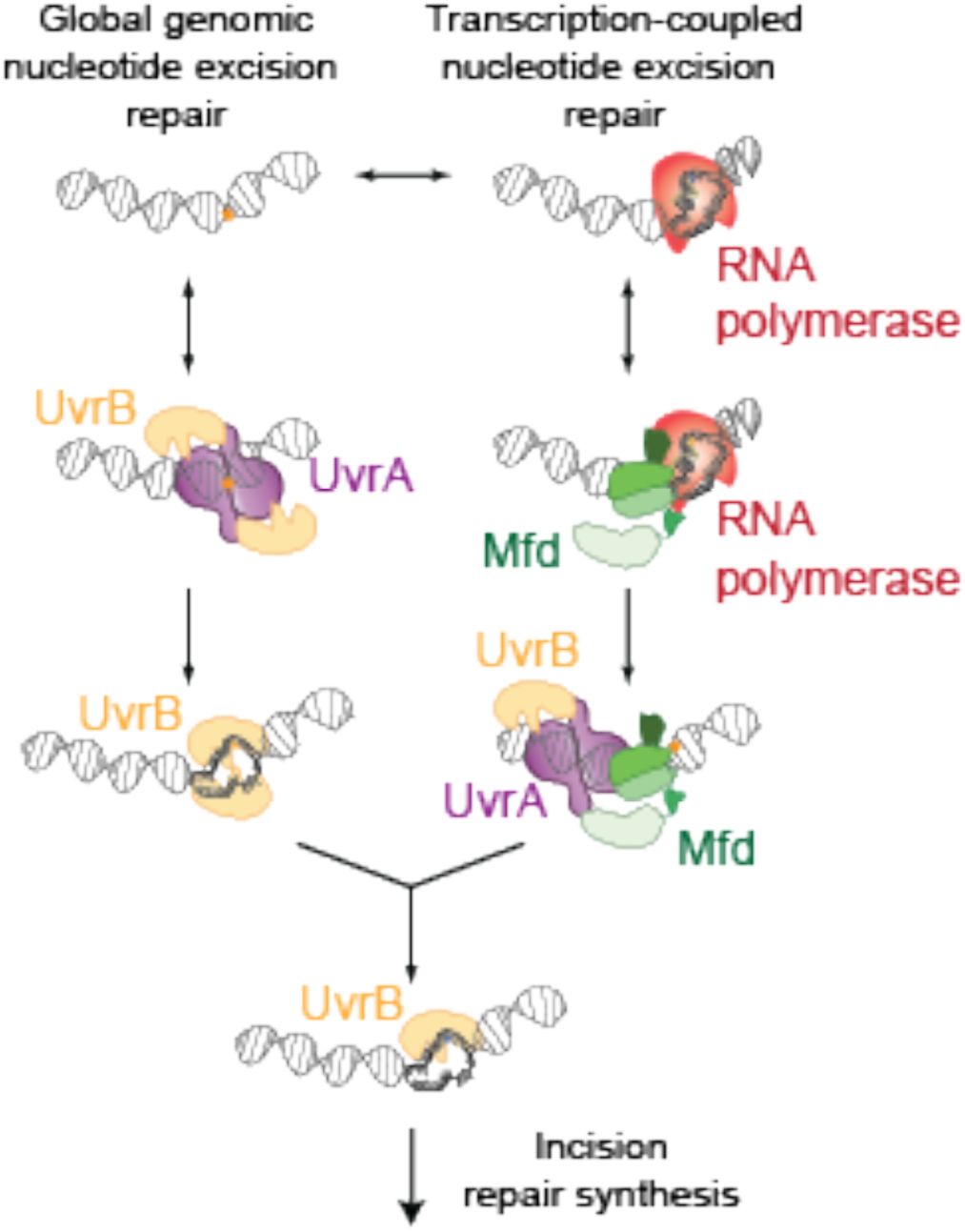
Nucleotide excision repair in *Escherichia coli*. Damage detection in nucleotide excision repair in *E. coli* proceeds *via* global damage surveillance executed by UvrA_2_(B), and RNA polymerase transcribing damaged template DNA. The UvrA dimer loads UvrB which verifies the presence of DNA damage in a strand-specific manner. Alternately, stalled elongation complexes at the site of DNA damage are rescued by the transcription repair coupling factor Mfd, which in turn recruits UvrA_2_(B) to the site of the stalled RNAP. This is followed by strand-specific loading of UvrB at the site of the lesion. Following damage verification by UvrB, a single-stranded patch of DNA containing the damage is incised by the UvrC endonuclease. This is followed by repair synthesis and ligation coordinated by UvrD, PolI and LigA.

In several studied organisms (barring certain archaea^4, 5^), removal of DNA damage also occurs via the triggering of strand-specific DNA repair following the stalling of RNA polymerase (RNAP) at sites of lesions (reviewed in ref.^6^). In this case, a ternary elongation complex (TEC) of RNAP that is unable to catalyse nascent RNA synthesis manifests as an ultra-stable protein-DNA roadblock. Transcription-repair coupling factors such as the prokaryotic Mfd, and the eukaryotic homologs Rad26/CSB are dedicated factors that recognize these TECs and remodel them^7, 8, 9, 10, 11^. In prokaryotes, Mfd is recruited to the site of a failed TEC, and in turn it recruits the UvrA_2_B complex via interactions between its UvrB-homology module (BHM) and UvrA. (Fig. 1) ^9, 11, 12, 13^. Similarly, in eukaryotes, CSB is recruited to the site of a stalled RNAPII complex, and recruits the TFIIH complex^14^.

DNA repair triggered by stalling of TECs is termed transcription-coupled repair (TCR), in contrast to the direct detection of lesions by the UvrAB damage sensor (global genomic repair, or GGR). Studies investigating the rate of repair during TCR *vs*. GGR, have reported an enhancement in the rate of removal of UV-induced lesions from the template strand in transcribed DNA compared to non-transcribed DNA^15, 16, 17, 18, 19, 20^. This observation has sparked several studies targeted at understanding the mechanistic basis of rate enhancement^13, 21, 22^. A recent single-molecule *in vitro* study reported that the time to incision in TCR is approximately three-fold faster than in GGR under certain conditions^13^.

A diverse set of intermediates is readily formed *in vitro -* ranging from a translocating RNAP-Mfd complex, arrested RNAP-Mfd-UvrA_2_ and the complete Mfd-UvrA_2_-UvrB handoff complex in the presence of both UvrA and UvrB^12, 13^. To understand which of these intermediates are formed inside cells, we have recently visualized Mfd in cells and quantified its lifetime in the TCR reaction^12^. A recent study failed to detect an influence of Mfd on the behaviour of fluorescently tagged UvrA in living cells^23^. Therefore, *in vitro* studies notwithstanding, how TCR is orchestrated by UvrA in cells remains unclear.

In this work, we revisited this question in the context of live cells and applied high-resolution single-molecule imaging methods that permit accurate measurements of DNA binding lifetimes over a broad timescale ranging from a few hundred milliseconds to several minutes^12, 24^. We asked the question: what is the lifetime of interactions of UvrA with Mfd and UvrB? To answer this question, we visualized fluorescently tagged UvrA in cells and measured the lifetimes of the interactions with DNA in wild-type and TCR-deficient cells. We find that UvrA is long lived on DNA in the absence of UvrB and Mfd, and that its dissociation is promoted by UvrB in cells. The cellular concentration of UvrA relative to UvrB strongly influences its binding lifetime in interactions with DNA-bound Mfd. The binding lifetimes of Mfd-UvrA interactions are only detected under conditions of limiting UvrB in the absence of exogenous DNA damage, suggesting a role for UvrB in resolving Mfd-UvrA intermediates. Exposure to ultraviolet light (UV) leads to an increase in the binding lifetime of UvrA in TCR-deficient cells. In contrast, in TCR-proficient cells, the DNA-bound lifetimes of UvrA and Mfd are identical, and drop upon UV exposure suggesting the formation of an Mfd-UvrA intermediate in TCR. Together, these studies characterize the interaction of UvrA with Mfd in live cells. Here, we establish a comprehensive framework for characterizing the binding kinetics of DNA repair proteins participating in multiple parallel pathways *in vivo* using non-perturbative, single-molecule imaging approaches.

## RESULTS

The nucleotide excision repair reaction can be considered to occur in five discrete stages: damage recognition, damage verification, incision, repair synthesis and ligation (reviewed in ref^2^). UvrA mediates damage recognition by itself and by interacting with Mfd^1^. It does so by surveying the genome constantly for the presence of DNA damage or for Mfd-bound TECs. Following DNA binding at a putative damage site, damage verification is then orchestrated by the damage verification enzyme UvrB^25^. Surprisingly, UvrA by itself has relatively poor specificity for binding to DNA damage over undamaged DNA^26^. Here, we set out to characterize the kinetic states that describe damage surveillance and loading of UvrB in live cells.

### Imaging of UvrA-YPet in the absence of exogenous DNA damage

To visualize the binding of UvrA to DNA in cells, we replaced the *uvrA* gene with a C-terminal fusion of *uvrA* to the gene for the yellow fluorescent protein (YPet^27^) in MG1655 cells using λ Red recombination (Fig. 2a)^12, 28^. This strategy enabled observation of fluorescent UvrA-YPet expressed from the native, SOS-inducible *uvrA* promoter (Supplementary Movie 1). We first performed UV-survival assays to assess the ability of UvrA-YPet to execute nucleotide excision repair (NER). Compared to wild-type cells, *uvrA-YPet* cells exhibited somewhat poorer survival upon exposure to UV (Supplementary Fig. 1a). Considering that C-terminal fusions of UvrA are fully functional in NER^23, 29^, this modestly lower survival of *uvrA-YPet* cells may be attributable to a lower efficiency of protein translation.

**Figure 2:**
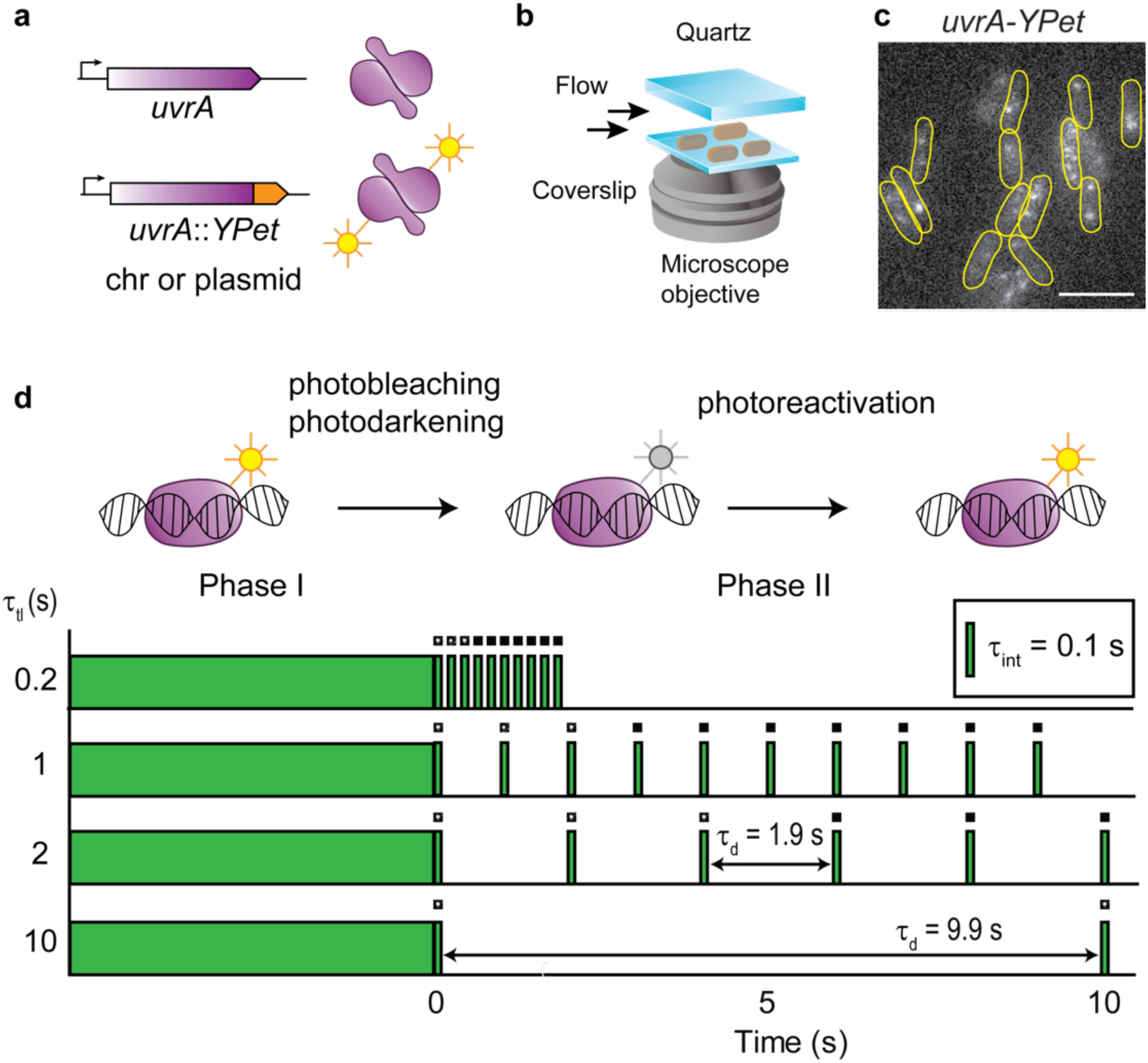
Construction of *uvrA-YPet* and imaging of UvrA-YPet. **a.** A chromosomal fusion of UvrA to the yellow fluorescent protein (YPet) under the native *uvrA* locus was created using λ Red recombination in MG1655 cells. In a second approach, the *uvrA-YPet* allele was expressed under the native *uvrA* promoter from a low-copy plasmid in Δ*uvrA* cells. **b.** Cells expressing fluorescent UvrA-YPet were grown to early exponential phase and loaded in a flow cell. Cells were imaged under constant supply of aerated growth medium for several hours. **c.** Fluorescence images of UvrA-YPet reveal a mixture of foci and diffuse cellular background signal. Scale bar represents 5 μm. Cell outlines are provided as a guide to the eye. **d.** Schematic of interval imaging approach employed to measure the off rates of fluorescently tagged proteins in cells. Each acquisition is collected in two phases. In the first phase, fluorescent signal is bleached to enable observation of single fluorescent YPet molecules. In the second phase, a dark frame τ_d_ is introduced such that the time-lapse time τ_tl_ = τ_d_ + τ_int_, where τ_int_ is the integration time (100 ms). In this phase, the lifetimes of individual binding events of UvrA-YPet molecules are measured and combined to obtain a cumulative residence time distribution.

Therefore, we set out to measure the copy numbers of UvrA-YPet in *uvrA-YPet* cells grown in EZ-rich defined media supplemented with glucose at 30 °C. Exponentially growing cells were deposited on a modified glass coverslip at the bottom of a flow cell and visualized by illumination with 514-nm laser light under continuous flow of growth medium (Fig. 2b). Images of *uvrA-YPet* cells revealed DNA-bound UvrA-YPet molecules that manifested as static foci and diffusive molecules contributing to cellular background fluorescence (Fig. 2c). These observations are consistent with its role as a damage surveillance protein.

Exposure to laser light led to rapid loss of YPet signal due to photodarkening and photobleaching of the chromophore (Supplementary Movie 1). We used this loss of signal to measure copy numbers of UvrA-YPet in cells. Dividing the background-corrected cellular fluorescence intensity by the intensity of a single YPet molecule revealed a copy number of 16 ± 4 copies of UvrA-YPet per cell (Supplementary Fig. 1b-d). Copy numbers of UvrA are strongly influenced by the carbon source present in the growth medium, ranging from 9 - 43 copies (minimal media) to 129 copies (rich media) per cell ^30^. These estimates exceed the copy numbers of UvrA-YPet detected in the *uvrA-YPet* strain grown in rich medium. The lower copy numbers of UvrA-YPet are consistent with the minor deficiencies in survival observed after UV exposure (Supplementary Fig. 1a).

To confirm the hypothesis that the deficiencies in UV survival observed for the *uvrA-YPet* strain are attributable to lower copy numbers as opposed to a catalytically deficient protein, we created a low copy plasmid (pSC101 origin of replication, 3-4 copies/cell^31^) expressing the C-terminal YPet fusion of UvrA under its native promoter (pUvrA-YPet). We then transformed Δ*uvrA* cells with pUvrA-YPet. These cells express UvrA-YPet to the extent of 120 ± 28 copies per cell (Supplementary Fig. 1b-d). UV survival assays (Supplementary Fig. 1e) revealed that UvrA-YPet expressed from the plasmid is able to fully complement the Δ*uvrA* phenotype. Therefore, we conclude that the copy numbers of UvrA-YPet expressed from the endogenous *uvrA* promoter represent lower copy numbers compared to untagged UvrA expressed in wild-type cells, likely reflecting a poorer efficiency of translation of the *uvrA-YPet* gene.

### Interval imaging strategy to measure DNA binding kinetics

Continuous imaging of UvrA-YPet could not be used to measure DNA binding lifetimes, since the apparent lifetime of a focus represents UvrA-YPet molecules dissociating from the site or bound UvrA-YPet molecules that are photobleached during the imaging. Consequently, measurement of interactions that last longer than the photobleaching lifetime is impossible. Instead we imaged UvrA-YPet using a strategy of performing time-lapse imaging with dark periods of varying intervals (for brevity, we term this mode of imaging ‘interval imaging’)^12, 24, 32, 33^ (Fig. 2d) that elegantly deconvolutes the lifetime of the interaction of UvrA-YPet with DNA and the lifetime of the fluorescent probe. Briefly, the introduction of a dark interval (τ_d_) between consecutive frames of duration (τ_int_) extends the observation time window. Here the time between consecutive frames is denoted as time-lapse time (τ_tl_). By acquiring the same number of frames in each video collected with a different dark interval, the photobleaching rate (*k*_b_) is maintained constant (see Table 1 and Methods) while the observation window is extended arbitrarily. From these videos, cumulative residence time distributions of DNA-bound UvrA-YPet are constructed. Since these distributions reflect a mixture of two populations (UvrA molecules that dissociate and YPet molecules that photobleached), fitting them to an exponential function yields an effective off rate (*k*_eff_)^32^. The product (*k*_eff_τ_tl_) is a linear function of true off rate (*k*_off_τ_tl_) and the normalized photobleaching rate (*k*_b_τ_int_)^32^. For purposes of illustration, the data can be represented as *k*_eff_τ_tl_ *vs.* τ_tl_ plots where the slope reveals the off rate *k*_off_ and the Y-intercept reveals *k*_b_. This interval imaging strategy enables accurate quantification of binding lifetimes over three orders of magnitude from 0.1 s to several minutes^12, 32, 34^.

**Table 1:**
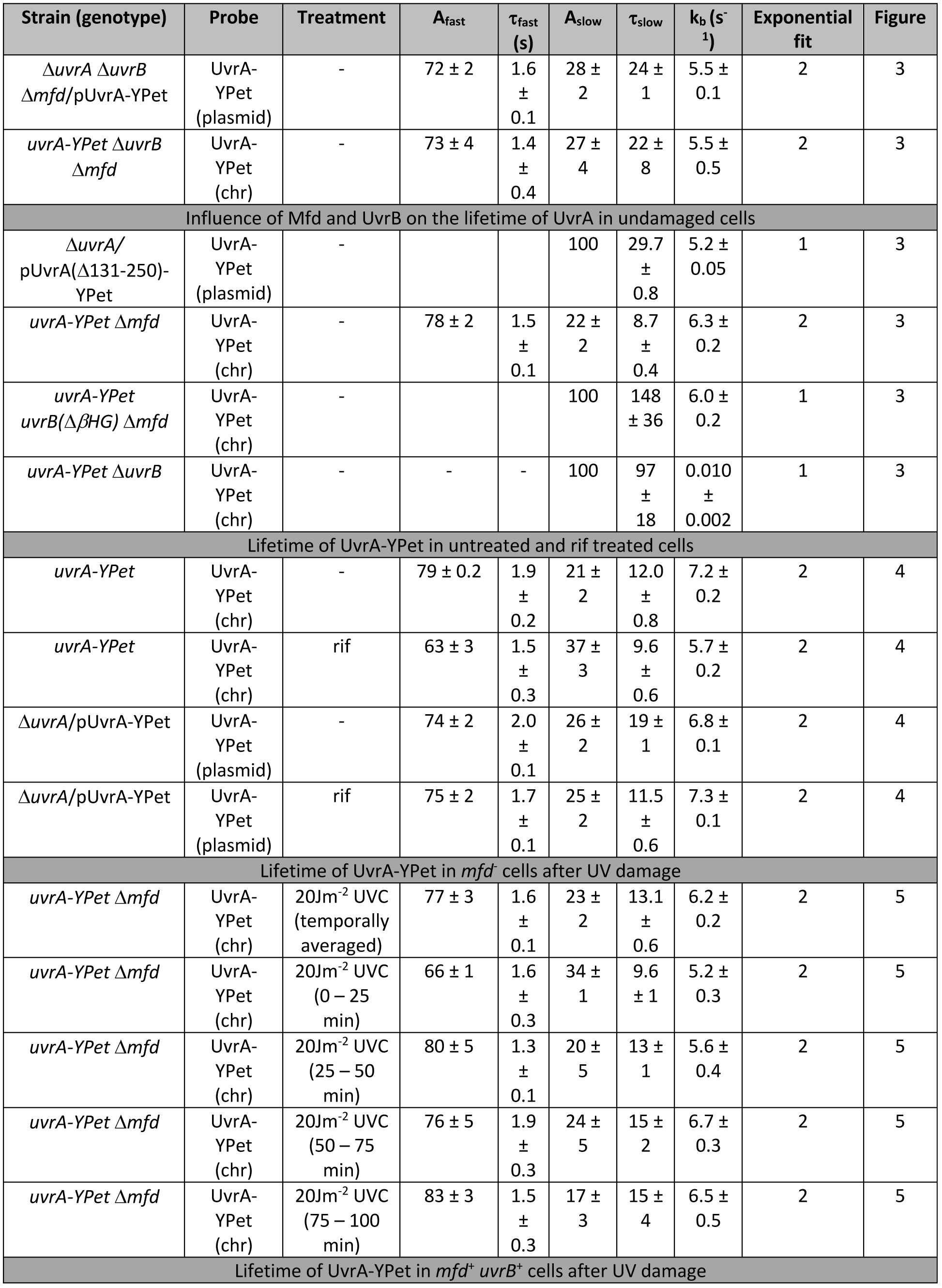
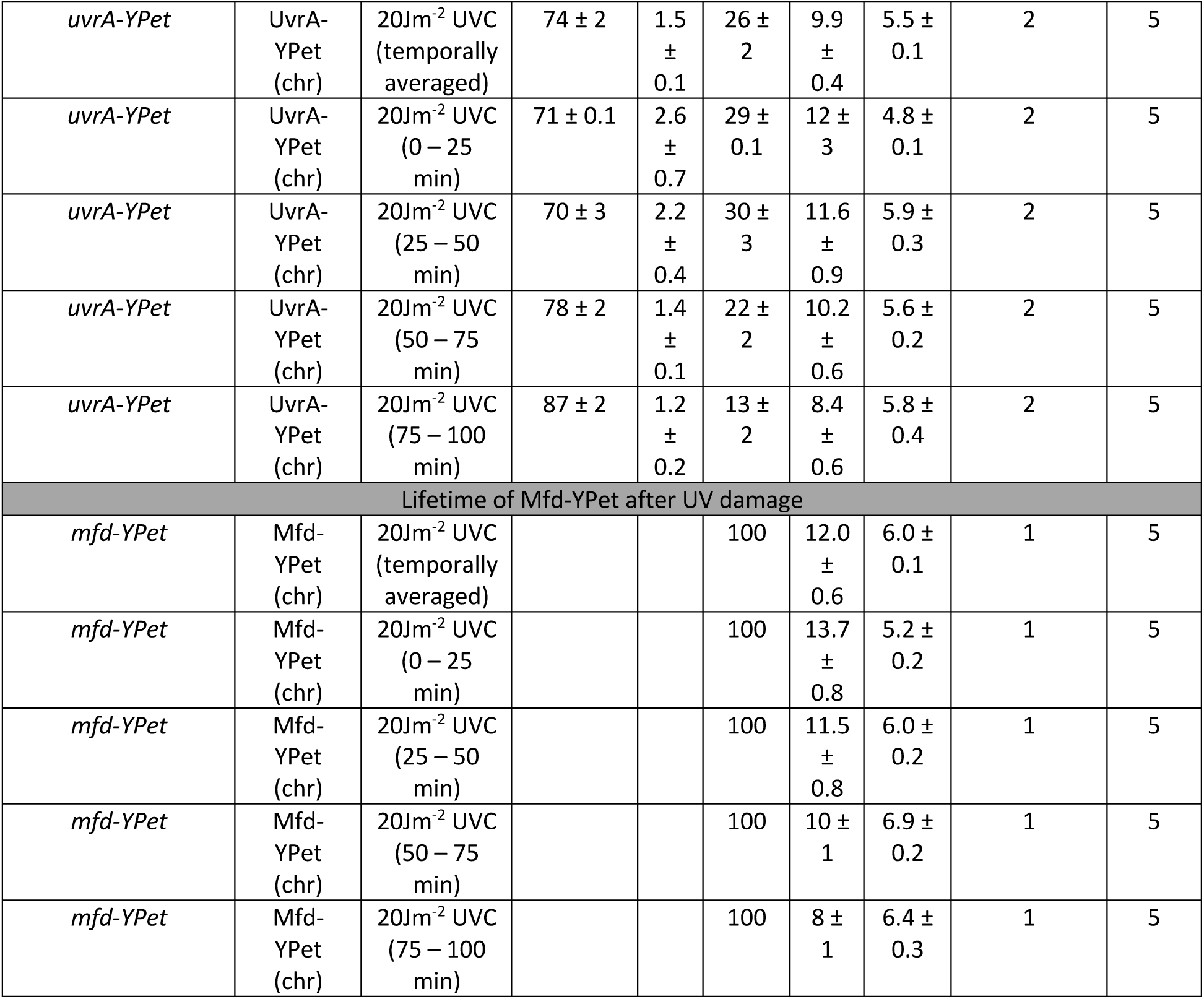
Summary of binding lifetimes.

| Strain (genotype) | Probe | Treatment | A<sub>fast</sub> | τ<sub>fast</sub> (s) | A<sub>slow</sub> | τ<sub>slow</sub> | k<sub>b</sub> (s<sup>-1</sup>) | Exponential fit | Figure |
| --- | --- | --- | --- | --- | --- | --- | --- | --- | --- |
| *ΔuvrA ΔuvrB Δmfd*/pUvrA-YPet | UvrA-YPet (plasmid) | - | 72 ± 2 | 1.6 ± 0.1 | 28 ± 2 | 24 ± 1 | 5.5 ± 0.1 | 2 | 3 |
| *uvrA-YPet ΔuvrB Δmfd* | UvrA-YPet (chr) | - | 73 ± 4 | 1.4 ± 0.4 | 27 ± 4 | 22 ± 8 | 5.5 ± 0.5 | 2 | 3 |
| Influence of Mfd and UvrB on the lifetime of UvrA in undamaged cells |  |  |  |  |  |  |  |  |  |
| *ΔuvrA*/pUvrA(Δ131-250)-YPet | UvrA-YPet (plasmid) | - |  |  | 100 | 29.7 ± 0.8 | 5.2 ± 0.05 | 1 | 3 |
| *uvrA-YPet Δmfd* | UvrA-YPet (chr) | - | 78 ± 2 | 1.5 ± 0.1 | 22 ± 2 | 8.7 ± 0.4 | 6.3 ± 0.2 | 2 | 3 |
| *uvrA-YPet uvrB(ΔβHG) Δmfd* | UvrA-YPet (chr) | - |  |  | 100 | 148 ± 36 | 6.0 ± 0.2 | 1 | 3 |
| *uvrA-YPet ΔuvrB* | UvrA-YPet (chr) | - | - | - | 100 | 97 ± 18 | 0.010 ± 0.002 | 1 | 3 |
| Lifetime of UvrA-YPet in untreated and rif treated cells |  |  |  |  |  |  |  |  |  |
| *uvrA-YPet* | UvrA-YPet (chr) | - | 79 ± 0.2 | 1.9 ± 0.2 | 21 ± 2 | 12.0 ± 0.8 | 7.2 ± 0.2 | 2 | 4 |
| *uvrA-YPet* | UvrA-YPet (chr) | rif | 63 ± 3 | 1.5 ± 0.3 | 37 ± 3 | 9.6 ± 0.6 | 5.7 ± 0.2 | 2 | 4 |
| *ΔuvrA*/pUvrA-YPet | UvrA-YPet (plasmid) | - | 74 ± 2 | 2.0 ± 0.1 | 26 ± 2 | 19 ± 1 | 6.8 ± 0.1 | 2 | 4 |
| *ΔuvrA*/pUvrA-YPet | UvrA-YPet (plasmid) | rif | 75 ± 2 | 1.7 ± 0.1 | 25 ± 2 | 11.5 ± 0.6 | 7.3 ± 0.1 | 2 | 4 |
| Lifetime of UvrA-YPet in *mfd*<sup>-</sup> cells after UV damage |  |  |  |  |  |  |  |  |  |
| *uvrA-YPet Δmfd* | UvrA-YPet (chr) | 20Jm<sup>-2</sup> UVC (temporally averaged) | 77 ± 3 | 1.6 ± 0.1 | 23 ± 2 | 13.1 ± 0.6 | 6.2 ± 0.2 | 2 | 5 |
| *uvrA-YPet Δmfd* | UvrA-YPet (chr) | 20Jm<sup>-2</sup> UVC (0 – 25 min) | 66 ± 1 | 1.6 ± 0.3 | 34 ± 1 | 9.6 ± 1 | 5.2 ± 0.3 | 2 | 5 |
| *uvrA-YPet Δmfd* | UvrA-YPet (chr) | 20Jm<sup>-2</sup> UVC (25 – 50 min) | 80 ± 5 | 1.3 ± 0.1 | 20 ± 5 | 13 ± 1 | 5.6 ± 0.4 | 2 | 5 |
| *uvrA-YPet Δmfd* | UvrA-YPet (chr) | 20Jm<sup>-2</sup> UVC (50 – 75 min) | 76 ± 5 | 1.9 ± 0.3 | 24 ± 5 | 15 ± 2 | 6.7 ± 0.3 | 2 | 5 |
| *uvrA-YPet Δmfd* | UvrA-YPet (chr) | 20Jm<sup>-2</sup> UVC (75 – 100 min) | 83 ± 3 | 1.5 ± 0.3 | 17 ± 3 | 15 ± 4 | 6.5 ± 0.5 | 2 | 5 |
| Lifetime of UvrA-YPet in *mfd*<sup>+</sup> *uvrB*<sup>+</sup> cells after UV damage |  |  |  |  |  |  |  |  |  |
| *uvrA-YPet* | UvrA-YPet (chr) | 20Jm<sup>-2</sup> UVC (temporally averaged) | 74 ± 2 | 1.5 ± 0.1 | 26 ± 2 | 9.9 ± 0.4 | 5.5 ± 0.1 | 2 | 5 |
| *uvrA-YPet* | UvrA-YPet (chr) | 20Jm<sup>-2</sup> UVC (0 – 25 min) | 71 ± 0.1 | 2.6 ± 0.7 | 29 ± 0.1 | 12 ± 3 | 4.8 ± 0.1 | 2 | 5 |
| *uvrA-YPet* | UvrA-YPet (chr) | 20Jm<sup>-2</sup> UVC (25 – 50 min) | 70 ± 3 | 2.2 ± 0.4 | 30 ± 3 | 11.6 ± 0.9 | 5.9 ± 0.3 | 2 | 5 |
| *uvrA-YPet* | UvrA-YPet (chr) | 20Jm<sup>-2</sup> UVC (50 – 75 min) | 78 ± 2 | 1.4 ± 0.1 | 22 ± 2 | 10.2 ± 0.6 | 5.6 ± 0.2 | 2 | 5 |
| *uvrA-YPet* | UvrA-YPet (chr) | 20Jm<sup>-2</sup> UVC (75 – 100 min) | 87 ± 2 | 1.2 ± 0.2 | 13 ± 2 | 8.4 ± 0.6 | 5.8 ± 0.4 | 2 | 5 |
| Lifetime of Mfd-YPet after UV damage |  |  |  |  |  |  |  |  |  |
| *mfd-YPet* | Mfd-YPet (chr) | 20Jm<sup>-2</sup> UVC (temporally averaged) |  |  | 100 | 12.0 ± 0.6 | 6.0 ± 0.1 | 1 | 5 |
| *mfd-YPet* | Mfd-YPet (chr) | 20Jm<sup>-2</sup> UVC (0 – 25 min) |  |  | 100 | 13.7 ± 0.8 | 5.2 ± 0.2 | 1 | 5 |
| *mfd-YPet* | Mfd-YPet (chr) | 20Jm<sup>-2</sup> UVC (25 – 50 min) |  |  | 100 | 11.5 ± 0.8 | 6.0 ± 0.2 | 1 | 5 |
| *mfd-YPet* | Mfd-YPet (chr) | 20Jm<sup>-2</sup> UVC (50 – 75 min) |  |  | 100 | 10 ± 1 | 6.9 ± 0.2 | 1 | 5 |
| *mfd-YPet* | Mfd-YPet (chr) | 20Jm<sup>-2</sup> UVC (75 – 100 min) |  |  | 100 | 8 ± 1 | 6.4 ± 0.3 | 1 | 5 |

### UvrA is long-lived on DNA in the absence of UvrB and Mfd

First, we interrogated UvrA binding kinetics in the absence of its two major interacting partners UvrB and Mfd in growing cells. To that end, we transformed cells lacking UvrA, UvrB and Mfd (Δ*uvrA* Δ*uvrB* Δ*mfd* cells) with pUvrA-YPet. In these cells, we expected that interactions of UvrA-YPet with chromosomal DNA would reflect two of its key activities: binding to non-damaged DNA and binding to endogenous DNA damage produced as a by-product of cellular metabolism (Fig. 3a). Indeed, measurements of UvrA-YPet kinetics of dissociation in these cells revealed two lifetimes that are an order of magnitude apart - a fast lifetime (τ_UvrA|Δ*uvrA* Δ*uvrB* Δ*mfd*, fast_) of 1.6 ± 0.1 s (72 ± 2 %) and a slow lifetime (τ_UvrA|Δ*uvrA* Δ*uvrB* Δ*mfd*, slow_) of 24 ± 1 s (28 ± 2 %) (summarized in Fig. 3c; Supplementary Figure 2a-b, and Table 1; error bars represent standard deviation of the bootstrap distribution of values obtained by performing global fits to CRTDs ten times). To eliminate the possibility that this measured lifetime is affected by cellular copy numbers of UvrA, we additionally created a strain that expresses *uvrA-YPet* from its endogenous promoter, and lacks the genes for *uvrB* and *mfd* (*uvrA-YPet* Δ*uvrB* Δ*mfd*). The measured lifetimes of UvrA-YPet in this strain were found to be identical within error (τ_UvrA|*uvrA-Ypet* Δ*uvrB* Δ*mfd*, slow_ = 22 ± 8 s (27 ± 4 %) and τ_UvrA|*uvrA-YPet* Δ*uvrB* Δ*mfd*, fast_ = 1.4 ± 0.4 s (73 ± 4%)) to those in the Δ*uvrA* Δ*uvrB* Δ*mfd*/pUvrA-YPet strain (Fig. 3a and 3c, Supplementary Figure 2c-d and Table 1). Further, we also measured the binding lifetime of a mutant UvrA that is deficient in its interactions with UvrB and Mfd (Fig. 3b). Since UvrA interacts with both UvrB and Mfd *via* the interface formed by residues 131-250 ^35, 36, 37^, we expected that the labelled mutant UvrA lacking residues 131-250, UvrA(Δ131-250)-YPet, would be a faithful reporter of binding of kinetics of UvrA alone in *uvrB*^+^ *mfd*^+^ cells (Fig. 3b). Previous biochemical characterization of a UvrA(Δ131-248) mutant revealed that this mutant is unable to interact with UvrB and catalyze TCR and GGR^22^. Indeed, interval imaging of UvrA(Δ131-250)-YPet expressed from a low-copy plasmid (pUvrA(Δ131-250)-YPet) in Δ*uvrA* cells produced a binding lifetime (τ_UvrA(Δ131−250_) of 29.7 ± 0.8 s (summarized in Fig. 3c, Supplementary Figure 2e-f, Table 1). It is noteworthy that deletion of these residues leads to a complete abolishment of the short-lived species. The reasons for this loss may lie in structural differences between the wild-type and mutant proteins. Together, these results demonstrate that UvrA-YPet by itself is long-lived on DNA. Notably, the lifetimes measured in our experiments reveal values that are larger than previous *in vitro* measurements of UvrA binding^29^. The presence of two binding lifetimes reveals the presence of two populations of UvrA on DNA.

**Figure 3:**
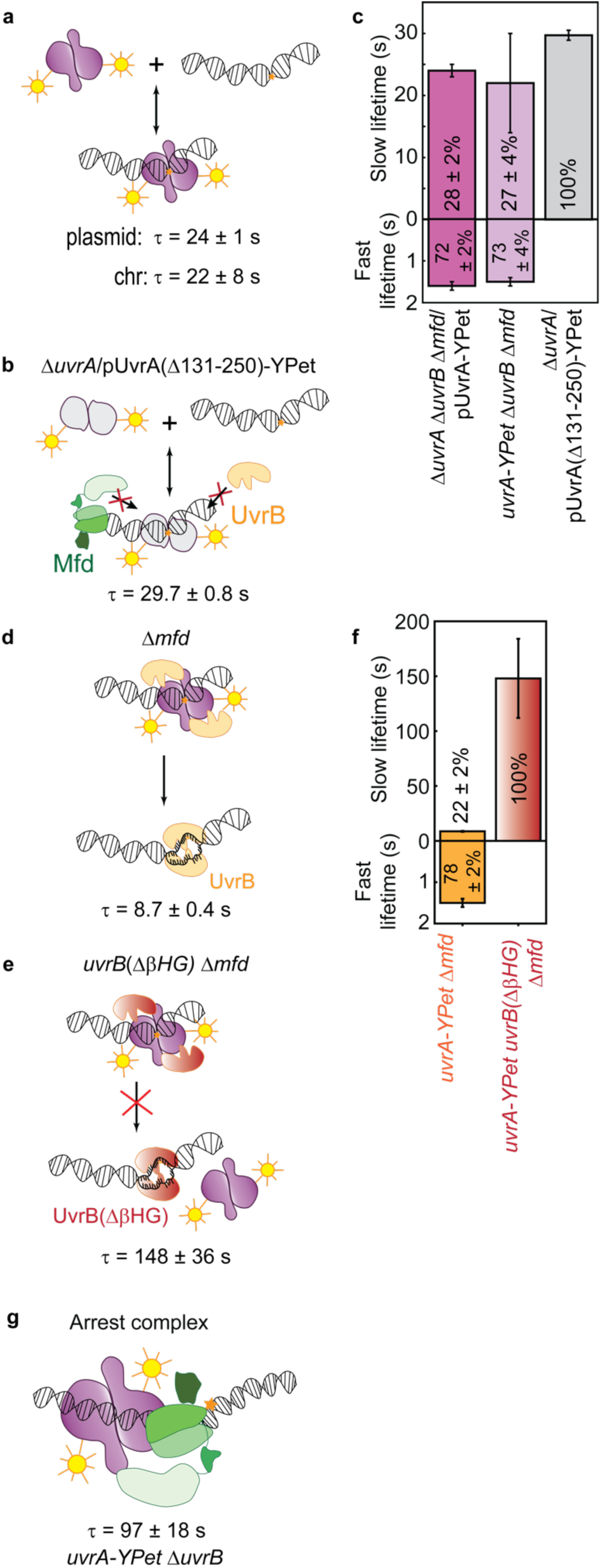
DNA binding lifetimes of UvrA-YPet in the presence of UvrB. **a.** Kinetics of UvrA-YPet interactions with DNA can be detected in the absence of UvrB and Mfd, in Δ*uvrA* Δ*uvrB* Δ*mfd* cells expressing UvrA-YPet from a low-copy plasmid or in Δ*uvrB* Δ*mfd* cells expressing UvrA-YPet from the chromosome. **b.** Cartoon illustrates DNA binding by the mutant UvrA(Δ131-250)-YPet, which is defective in interacting with UvrB and Mfd. **c.** Bar plots represent lifetimes of DNA-bound UvrA-YPet (plasmid: n = 34,927 counts from 5 repeats; chromosomal: n = 970r counts from 3 repeats) and mutant UvrA(Δ131-250)-YPet (a total of n = 88,232 counts from 4 repeats) in the corresponding genetic background. Lifetimes were obtained from globally fitting the cumulative residence time distributions (CRTDs) (see also Supplementary Fig. 2a-f). Where two kinetic sub-populations are detected, the fast lifetime is displayed in the lower panel. Percentage represents the amplitude of kinetic sub-populations. Error bars are standard deviations from ten bootstrapped CRTDs. See also source data file. **d.** Cartoon illustrates loading of UvrB by UvrA. This may occur at sites of undamaged or damaged DNA. **e.** Cartoon illustrates the complex formed by UvrA and the mutant UvrB(ΔβHG) that is deficient in loading reaction. **f.** Bar plots represent lifetimes of DNA-bound UvrA-YPet in Δ*mfd* cells expressing either wild-type UvrB (n = 29,743 counts from 11 repeats) or mutant UvrB(ΔβHG) (n = 16,353 counts from 2 repeats). Lifetimes were obtained from globally fitting the CRTDs, with more than 1,000 counts each CRTD (see Supplementary Fig. 2g - j). Where two kinetic sub-populations are detected, the fast lifetime is displayed in the lower panel. Percentage represents the amplitude of kinetic sub-populations. Error bars are standard deviations from ten bootstrapped CRTDs. See also source data file. **g.** Cartoon of the arrested Mfd-UvrA complex. See also Supplementary Figure 3 a-b.

### UvrB dissociates DNA-bound UvrA in cells lacking Mfd

Next, we studied the influence of UvrB on the DNA binding lifetime of UvrA in cells lacking Mfd (*uvrA-YPet* Δ*mfd* cells; Fig. 3d) during normal growth. In these cells, UvrA-YPet dissociated with a fast lifetime (τ_UvrA| Δ*mfd*, fast_) of 1.5 ± 0.1 s (amplitude: 78 ± 2%) and a slow lifetime (τ_UvrA| Δ*mfd*, slow_) of 8.7 ± 0.4 s (22 ± 2%) (summarized in Fig. 3f; Supplementary Figure 2g-h, Table 1). The lifetime of the slowly dissociating species observed in our measurements matches the lifetime detected for the dissociation of UvrA in the presence of UvrB previously (7 s) ^29^, and is consistent with measurements of UvrB loading at sites of DNA damage^38^. Notably, the fast lifetime is consistent with measurements from a previous study^23^; however, in this study a long-lived population of UvrA was not detected in the absence of exogenous DNA damage^23^.

Several *in vitro* studies have revealed that damage detection during NER proceeds *via* the loading of UvrB on DNA, followed by damage verification mediated *via* the helicase activity of UvrB ^39, 40, 41^. To investigate whether the ability of UvrB to be loaded on DNA *via* its β-hairpin is essential for dissociation of UvrA from DNA, we measured the lifetime of UvrA-YPet in *uvrA-YPet* Δ*mfd* cells expressing the β-hairpin deletion mutant of UvrB from the native *uvrB* locus (Methods). This mutant, UvrB(ΔβHG), is inefficiently loaded on DNA *in vitro* ^42^. The lifetime (τ_UvrA| *uvrB*(Δβ*HG*) Δ*mfd*_) of UvrA-YPet in cells expressing UvrB(ΔβHG) from the chromosome was 148 ± 36 s (100%); over 15-fold longer than that of UvrA-Ypet in cells lacking Mfd (Fig. 3e-f, Supplementary Figure 2i-j and Table 1). These data indicate that the UvrA-UvrB(ΔβHG) complex is arrested on DNA.

The lack of a short-lived species of UvrA in cells expressing mutant UvrB implies that the detectable population of UvrA can be sequestered to the chromosome in the form of a long-lived complex. Such a complex has been detected in single-molecule DNA stretching assays where UvrAB was demonstrated to slide on DNA ^29^. Additionally, we infer that in wild-type cells, loading of UvrB on DNA must promote the dissociation of UvrA. Indeed, our single-molecule live-cell imaging results highlight the physiological relevance of models constructed from *in vitro* studies that demonstrate that UvrB facilitates the dissociation of UvrA from DNA ^42, 43, 44^. These findings lead us to suggest that the 8.7s lifetime measured here corresponds to the lifetime of UvrA engaged in damage surveillance activities, where UvrA is turned over by UvrB loading.

### Mfd arrests UvrA on DNA in UvrB deficient cells

Next, we measured the lifetime of UvrA in cells lacking UvrB in the absence of exogenous DNA damage. Under these conditions, UvrA can form surveillance complexes (UvrA_2_) and interact with Mfd engaged with stalled RNAP. Interval imaging of UvrA-YPet in cells lacking UvrB (*uvrA-YPet* Δ*uvrB*) revealed a single long-lived UvrA species with a lifetime (τ_UvrA| Δ*uvrB*_) of 97 ± 18 s (Fig. 3g, Supplementary Figure 3a-b). Since UvrA alone binds DNA with a lifetime of approximately 24 s, this highly stable species must reflect interactions with Mfd. We propose that this slowly dissociating species reflects the arrested Mfd-UvrA complex^13^ in cells lacking UvrB. Our attempts at co-localization of UvrA and Mfd using the spectrally separated probes, YPet and PAmCherry, were limited by the poor expression of Mfd-PAmCherry and UvrA-PAmCherry in cells under our standard growth conditions (Supplementary Note 1 and Supplementary Figure 3c-d). Considering the copy numbers of UvrA measured here (16 per cell) and Mfd (22 per cell)^12^, and the reported picomolar affinity of UvrA for Mfd^13^, a large fraction of entire population of UvrA can be efficiently sequestered in complex with Mfd, consistent with the observation of only a single, slowly dissociating species in cells lacking *uvrB*.

### UvrA is longer lived on DNA in *mfd*^+^ cells compared to mfd^-^ cells in the absence of exogenous damage

We next set out to measure the residence time of DNA-bound UvrA in cells carrying both Mfd and UvrB in the absence of exogenous DNA damage. UvrA is recruited to DNA *via* Mfd to form the asymmetric handoff complex Mfd-UvrA_2_-UvrB that couples failed TECs to the repair machinery, unlike the symmetric UvrB-UvrA_2_-UvrB complex formed during damage surveillance in the absence of Mfd (Fig. 1). Our previous characterization of Mfd demonstrated that the residence time of Mfd is governed by UvrA, suggesting that UvrA is recruited to Mfd during normal growth^12^. Hence, we anticipated three scenarios for the lifetime of UvrA in wild-type cells. First, if the residence time of UvrA-YPet is equal to 8.7 s, this would indicate either that the lifetimes of Mfd-UvrA and UvrA-UvrB interactions are similar, or that the lifetimes are different but the recruitment of UvrAB to RNAP-bound Mfd occurs so infrequently that only UvrAB complexes are detected. Second, a lifetime shorter than 8.7 s would suggest that Mfd promotes the dissociation of UvrA. Finally, a lifetime longer than 8.7 s would indicate that Mfd stabilizes UvrA.

To distinguish between these three scenarios, we imaged UvrA-YPet in wild-type cells. Interval imaging of UvrA-YPet revealed a short-lived species with a lifetime (τ_UvrA, fast_) of 1.9 ± 0.2 s (79 ± 0.2%) and a long-lived species of UvrA with a lifetime (τ_UvrA, slow_) of 12.0 ± 0.8 s (21 ± 2%) (Fig. 4a, b, Supplementary Figure 4a-b, Table 1). To identify whether this lifetime reflected Mfd-dependent or independent interactions, we imaged *uvrA-YPet* cells in conditions where Mfd-RNAP interactions are abolished by rifampicin (rif) treatment. Interval imaging of UvrA-YPet in rif-treated cells revealed a short-lived species with a lifetime (τ_UvrA|rif,fast_) of 1.5 ± 0.3 s (63 ± 3 %) and a long-lived species of UvrA-YPet possessing a lifetime (τ_UvrA|rif, slow_) of 9.6 ± 0.6 s (37 ± 3 %) (Fig. 4a, c, Supplementary Figure 4c-d, Table 1). As expected, the slow lifetime of UvrA-YPet in rif-treated cells matches that in cells lacking Mfd. Further, measurements of UvrA-YPet fluorescence in rif-treated cells did not reveal a loss of YPet fluorescence arising from enhanced degradation of UvrA-YPet upon rif treatment (Supplementary Figure 4i). The decrease in the long-lived lifetime from 12.0 s to 9.6 s indicates that a slowly dissociating species (lifetime greater than 8.7 s) of UvrA is lost upon rif treatment, one that is involved in Mfd-dependent interactions. These results lead us to conclude that the interactions of Mfd-UvrAB complexes are more stable than UvrAB complexes alone.

**Figure 4:**
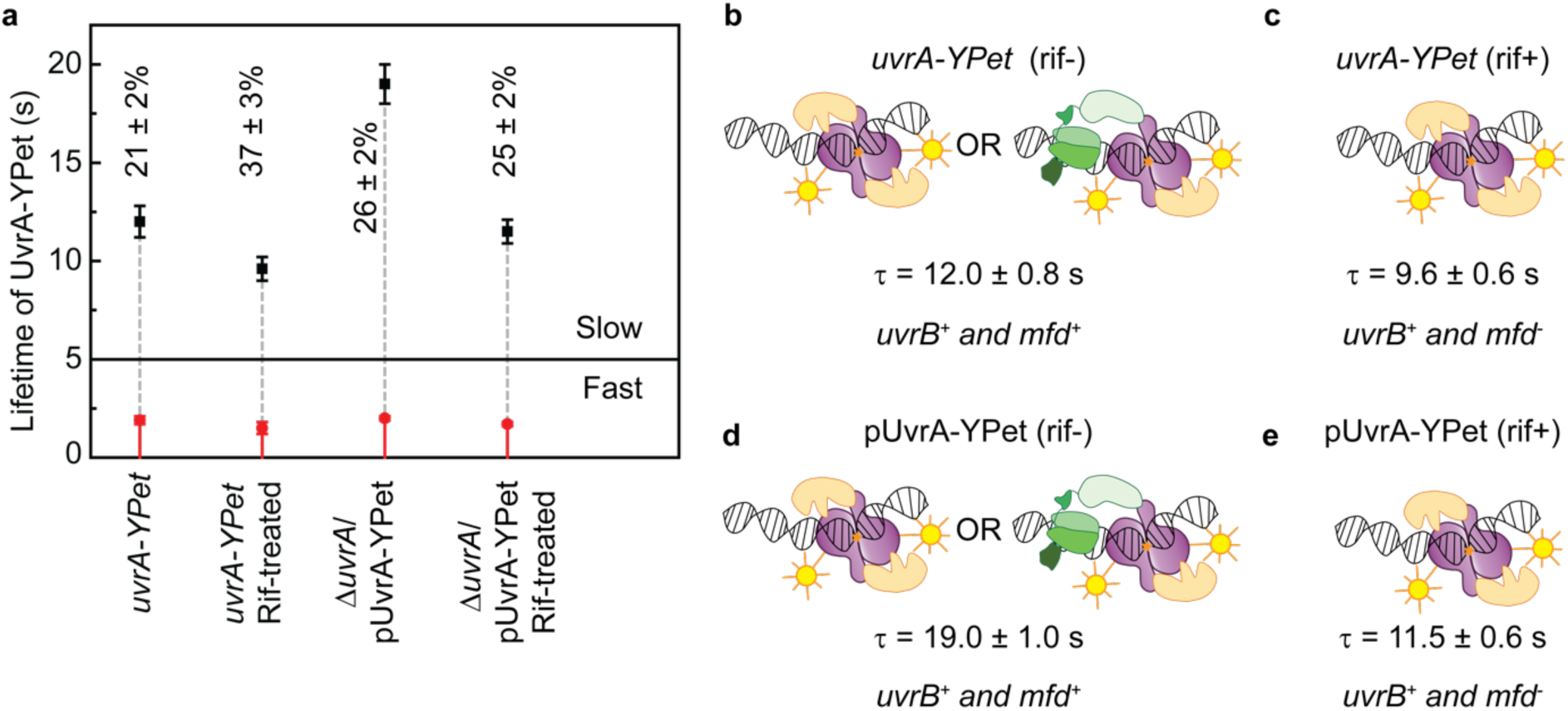
Dissociation kinetics of UvrA-YPet in TCR-competent cells. **a.** Lifetimes of UvrA-YPet in *uvrA-YPet* cells or Δ*uvrA*/pUvrA-YPet cells untreated or treated with rifampicin. Lifetimes were obtained from globally fitting the CRTDs (see Supplementary Figure 4 a, c, e, g). Slow lifetime is presented in black. Fast lifetime is presented in red. Percentages represent the amplitude of the slowly dissociating population. Error bars are standard deviations from ten bootstrapped CRTDs. See also source data file. **b.** In the presence of UvrB and Mfd, UvrA-YPet in *uvrA-YPet* cells exhibited a slow lifetime of 12.0 ± 0.8 s, reflecting UvrA interactions with UvrB and Mfd (n = 20,111 counts from 8 repeats). **c.** Rifampicin treatment abolishes Mfd-RNAP interactions, hence, UvrA-YPet is channelled towards interactions with UvrB, with the slow lifetime was found to be 9.6 ± 0.6 s (n = 15,355 counts from 3 repeats). **d.** When UvrA-YPet concentration increased eight fold via the means of plasmid expression, the slow lifetime of UvrA-YPet in Δ*uvrA*/pUvrA-YPet cells was found to be 19 ± 1 s, longer than that of UvrA-YPet in *uvrA-YPet* cells (12 s) (n = 19,853 counts from 3 repeats). **e.** Upon rifampicin treatment, the slow lifetime of UvrA-YPet in Δ*uvrA*/pUvrA-YPet cells reduced to 11.5 ± 0.6 s (n = 31,788 counts from 4 repeats).

Since no mutants of UvrA have been identified that exclusively form the Mfd-UvrA_2_-UvrB complex but not the UvrA_2_UvrB_2_ complex in cells, measurement of the lifetime of UvrA in the Mfd-UvrA_2_-UvrB complex is difficult. Nevertheless, it is conceivable that the lifetime of the Mfd-UvrA_2_-UvrB complex could be influenced by the availability of downstream factors including UvrAB. We tested this hypothesis by measuring the lifetime of UvrA-YPet in wild-type cells expressing UvrA-YPet from the plasmid (Δ*uvrA*/pUvrA-YPet). Since UvrA can engage UvrB in solution, we reasoned that higher expression levels of UvrA would bind UvrB to form the UvrA_2_B_2_ surveillance complex, leaving little free UvrB for engagement with Mfd-UvrA_2_ complexes. Consistent with this expectation, the data revealed a short-lived species with a lifetime (τ_UvrA|↑, fast_) of 2.0 ± 0.1 s (74 ± 2 %) and a previously unencountered population of long-lived UvrA possessing a lifetime (τ_UvrA|↑, slow_) of 19 ± 1 s (26 ± 2%) (Fig. 4a, d, Supplementary Figure 4e-f, Table 1).

In cells, UvrA is involved in target search (1.6 ± 0.1 s and 24 ± 1 s lifetimes, Fig. 3c) and damage surveillance as part of UvrA_2_B_2_ (8.7 s lifetime, Fig. 3f) in addition to Mfd-dependent UvrA(B) complexes (with lifetime of at least 12 s). To identify whether this long-lived UvrA species (19 ± 1 s lifetime) interacts with Mfd, we treated Δ*uvrA*/pUvrA-YPet cells with rifampicin. Under this condition, we expected to recover the lifetime of UvrA as part of UvrA_2_ or UvrA_2_B_2_ complexes. Indeed, measurements of lifetimes of UvrA-YPet in rif-treated Δ*uvrA*/pUvrA-YPet cells revealed a lifetime (τ_UvrA|↑rif, slow_) of 11.5 ± 0.6 s (25 ± 2%) and a short lifetime (τ_UvrA|↑rif, fast_) of 1.7 ± 0.1 s (75 ± 2%) (Fig. 4a, e, Supplementary Figure 4g-h, Table 1). The faster turnover of UvrA in response to rif treatment is consistent with the inference that the lifetime of UvrA in the Mfd-UvrA_2_-UvrB intermediate is longer than that in the UvrA_2_B_2_ intermediate in the absence of exogenous damage.

Notably, rif treatment of Δ*uvrA*/pUvrA-YPet cells yielded a lifetime (11.5 s) that is longer than that measured for rif-treated *uvrA-YPet* cells (9.6 s), and cells lacking *mfd* (8.7 s). The simplest explanation consistent with these observations is that under conditions of high relative UvrA/UvrB abundance the population is composed of UvrA_2_B_(2)_ complexes (lifetime of 8.7 s) and DNA-bound UvrA_2_ awaiting turnover by UvrB (lifetime of 24 s). We suggest that at higher cellular concentrations of UvrA relative to UvrB and Mfd, the existing population of UvrB is now required to turnover a greater number of UvrA molecules on undamaged DNA and at sites of endogenous damage, in addition to TCR intermediates. This model predicts that Mfd and UvrA must form a TCR intermediate whose disassembly is contingent on the arrival of UvrB. Together these results highlight two important features of prokaryotic NER: first, that damage surveillance by UvrA is highly dynamic and can be readily diverted from one pathway to the other depending on the condition, and second that the lifetime of individual actors is determined by the presence of downstream factors.

### UvrA-Mfd interactions are prioritized in cells after UV irradiation

Next, we set out to characterize the behaviour of the UvrA in response to DNA damage. UV irradiation leads to the formation of UV-induced lesions in the chromosome^45^. These in turn elicit the induction of the SOS response during which the expression of UvrA and UvrB are upregulated^46^. The elevated levels of UvrA promote rapid removal of UV lesions from the DNA. To quantify this SOS-induced upregulation in real time, we set out to monitor the relative abundance of UvrA in cells following UV irradiation.

Time-lapse experiments on UV-irradiated *uvrA-YPet* allowed us to monitor the cellular fluorescence of tagged UvrA as a function of time. We immobilized *uvrA-YPet* cells in a flow cell with a quartz window and delivered a dose of 20 Jm^-2^ of damaging 254-nm UV light (Fig. 5a). This was followed by acquiring a single snapshot upon laser illumination with 514-nm light, every five minutes for three hours. Quantification of cellular fluorescence intensities revealed that the integrated fluorescence intensities of single *uvrA-YPet* cells increase 30 minutes after UV exposure by three-fold, consistent with the rapid deregulation of the SOS inducible *uvrA* promoter (Fig. 5b)^47, 48^. UvrA copy numbers have been suggested to increase from 25 to 250 copies per cell after SOS induction^49^. We note that since the experimental conditions associated with these measurements are not available in the published literature, we are unable to effectively compare our measurements with these numbers. Nevertheless, the basal levels and the exact extent of fold-induction after SOS induction may depend on the nature and dosage of the genotoxin, as well as growth conditions such as medium and temperature. Consistent with this argument, the copy numbers of UvrA were found to rise six-fold within 40 minutes of UV exposure (40 Jm^-2^) in cells growing at 37 °C in previous work ^50^.

**Figure 5:**
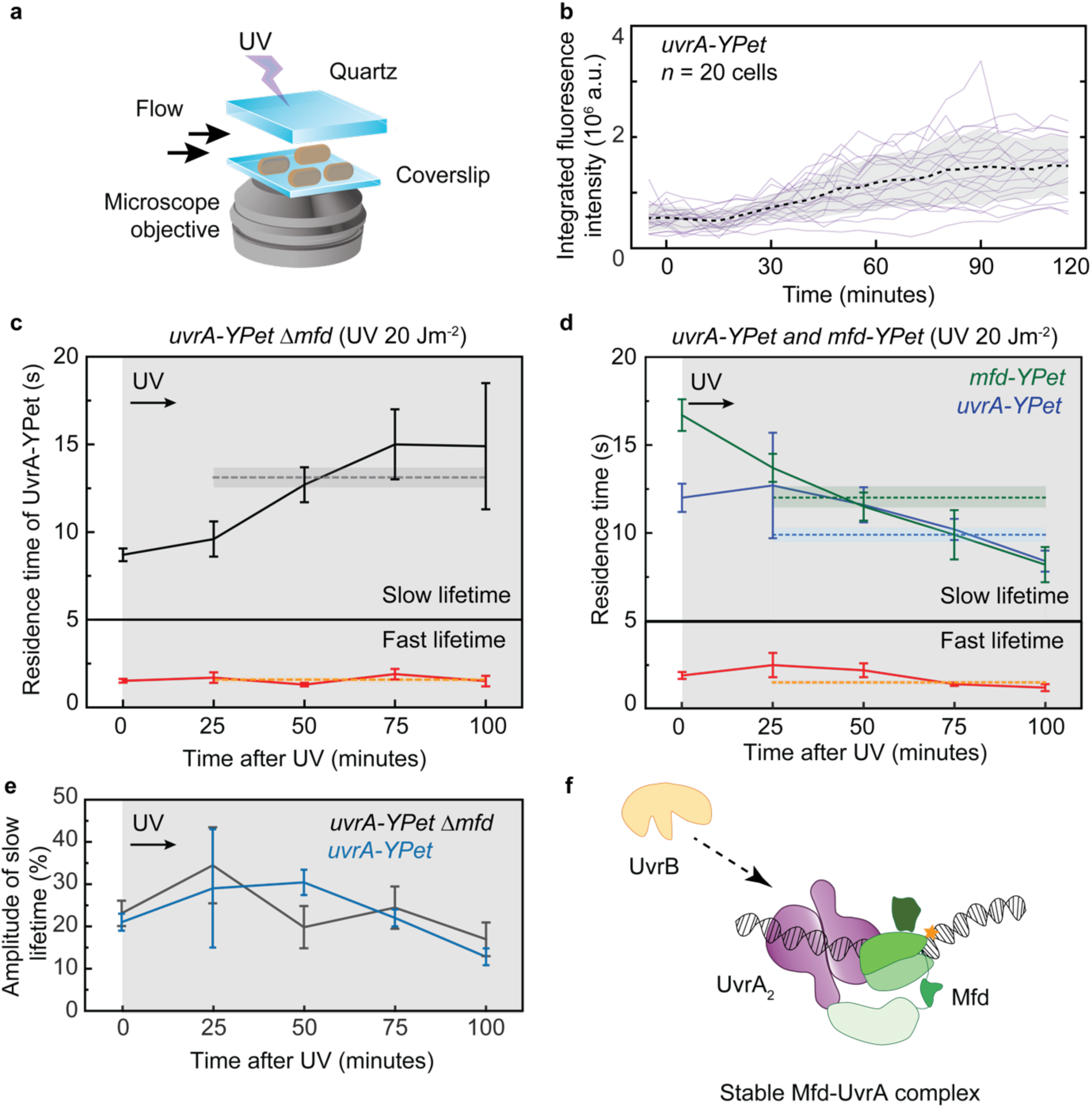
Lifetimes of DNA-bound UvrA and Mfd in response to UV irradiation. **a.** Experimental setup for introducing UV damage followed by interval imaging. A 20 Jm^-2^ dose of UV (254 nm) light is provided *via* a quartz window in the flow cell. This setup enables imaging of cells at 30 °C for several hours after UV. **b.** Fluorescence intensity of single *uvrA-YPet* cells increases following exposure to UV light, indicating that expression of UvrA-YPet is upregulated in the SOS response. **c.** Lifetimes of UvrA-YPet in TCR-deficient cells as a function of time following UV exposure. Lifetimes were obtained from globally fitting the CRTDs (see Supplementary Fig. 5a-d). Error bars are standard deviations from ten bootstrapped CRTDs. Lifetimes of the fast and slowly dissociating sub-populations are shown in the lower and upper panels respectively. Dashed lines represent lifetimes obtained from aggregated CRTDs within 100 minutes following UV exposure. The lifetimes at t = 0 minutes represent those of UvrA-YPet in TCR-deficient cells during normal growth and are reproduced from Fig. 4a. (After UV irradiation *uvrA-YPet* Δ*mfd* all data: n = 21,824 counts from 4 repeats; 0 – 25 min: n = 3243; 25 −50 min: n = 5175; 50 −75 min: n = 6079; 75 – 100 min: n = 5999 counts). **d.** Lifetimes of UvrA-YPet in *uvrA-YPet* cells (blue) (After UV irradiation *uvrA-YPet* all data: n = 25,935 counts from 4 repeats; 0 – 25 min: n = 5359; 25 −50 min: n = 8070; 50 −75 min: n = 7481; 75 – 100 min: n = 4983 counts) or Mfd-YPet in *mfd-YPet* cells (green) (After UV irradiation *mfd-YPet* all data: n = 14,553 counts from 4 repeats; 0 – 25 min: n = 5133; 25 −50 min: n = 3846; 50 −75 min: n = 3119; 75 – 100 min: n = 2286 counts) as a function of time following UV exposure. Lifetimes were obtained from globally fitting the CRTDs (see Supplementary Fig. 5a-d and Supplementary Fig. 6a-d). Error bars are standard deviations from ten bootstrapped CRTDs. Dash lines and error bands represent lifetimes and the corresponding standard deviations obtained from aggregated CRTDs within 100 minutes following UV exposure. Fitting results are available in Table 1. The lifetimes at t = 0 minutes represent those of UvrA-YPet in *uvrA-YPet* cells during normal growth and are reproduced from Fig. 4a. Lifetime of Mfd-YPet at t = 0 minutes is reproduced from our previous work (ref 12). **e.** Amplitudes of slowly dissociating species of UvrA-YPet in *mfd*^+^ (blue) or Δ*mfd* (black) cells carrying *uvrA-YPet.* See also table 1 for fitting results and source data file. **f.** UvrB (orange) controls the release of UvrA (purple) from UvrA-Mfd (green) intermediates.

Since UV-induced lesions are a substrate for UvrAB (reviewed in ref. ^1^), we then set out to measure the lifetime of UvrA in UV-irradiated cells. To that end, we irradiated TCR-deficient cells (*uvrA-Ypet* Δ*mfd*) with a pulse of 254-nm UV light delivered *in situ*. This was followed by interval imaging in four rounds, each lasting 25 minutes. Analysis of the complete data set revealed binding kinetics of UvrA-YPet corresponding to a short-lived species with a lifetime (τ_UvrA| Δ*mfd*, UV fast_) of 1.6 ± 0.1 s (77 ± 3%) and a long-lived species of UvrA corresponding to a lifetime of (τ_UvrA| Δ*mfd*, UV slow_) 13.1 ± 0.6 s (23 ± 2%) (Fig. 5c, and Supplementary Figure 5, Table 1). Strikingly the lifetime of the slowly dissociating species was larger than that detected in the absence of exogeneous DNA damage (8.7 s).

Seeking an explanation for the increase in binding lifetime of UvrA-YPet following UV exposure, we wondered if the longer lifetime of UvrA detected in these experiments represented temporally averaged measurements. Since each set of interval measurements lasted 25 min, we proceeded to disaggregate each data set into the four constitutive 25-min intervals after UV exposure. Analysis of the resulting data from each time window revealed that the measured lifetime of UvrA changes as a function of the experimental timeline after the UV pulse (Fig. 5c, and Supplementary Fig. 5, Table 1). Indeed, in the first 25 minutes, the slow lifetime of UvrA (9.6 ± 1 s) matched that measured in the absence of DNA damage (8.7 ± 0.4 s). This lifetime increased to a maximum of 15 ± 2 s in the 50-75 minute time window, finally plateauing to 15 ± 4 s in the 75-100 minute time window after UV exposure.

There are two main takeaways from these experiments. First, the lifetime of short-lived UvrA does not change upon UV exposure, and is identical to that measured in the absence of any exogenous DNA damage. We therefore conclude that this species is involved in binding undamaged DNA. Second, since the lifetime of long-lived UvrA changes upon UV exposure, we conclude that this species is engaged in DNA damage recognition.

Next, we repeated the interval imaging experiments on wild-type cells (*uvrA-YPet*) following exposure to a 20 Jm^-2^ pulse of 254-nm UV light provided *in situ*. In this case, UvrA-YPet exhibited two kinetic populations after UV-exposure, a short-lived population with a lifetime of 1.5 ± 0.1 s (74 ± 2 %) and a second, longer lived population with a lifetime of 9.9 ± 0.4 s (26 ± 2%) (Fig. 5d, and Supplementary Figure 6, Table 1). As before, we disaggregated each data set into the four constitutive 25-min intervals after UV exposure. In contrast to TCR-deficient cells, the measured lifetime of UvrA in wild-type cells remained low (8.4 ± 0.6 s) in the 75-100 minute time window after UV exposure. These data indicate that UvrA is turned over faster in an Mfd-dependent manner during the SOS response.

Next, we plotted the amplitude of the long lived species in each time-window after UV irradiation (Figure 5e). In *uvrA-YPet* cells, the fraction of the population dissociating with a slow off rate was greatest in the first 50 minutes after UV exposure (at approximately 30%), and subsequently dropped rapidly to 13 ± 2% at 100 minutes after UV. In comparison, *uvrA-YPet* Δ*mfd* cells exhibited a non-linear trajectory that nevertheless exhibited a similar reduction in the fraction of UvrA-YPet possessing a slow off rate. The most striking difference between the two strains was apparent at the 50 minute time point – cells carrying *mfd* exhibited a 1.5-fold greater fraction of UvrA molecules dissociating with the slow lifetime. These results are broadly consistent with the expectation that as repair progresses, fewer unrepaired lesions are available for UvrA to bind to.

We followed these studies with an investigation of the binding lifetimes of Mfd. In the absence of exogenous DNA damage, the lifetime of Mfd is approximately 18 s^12^. Interval imaging of Mfd-YPet in *mfd-YPet* cells exposed to UV light revealed that the lifetime of Mfd-YPet dropped during the course of the SOS response, leading to an average lifetime (τ_Mfd|UV_) of 12 s (Fig. 5d, and Supplementary Figure 7, Table 1). Strikingly, the binding lifetime of Mfd mirrored that of UvrA in wild-type cells in the time window from 25-100 min after UV, providing further evidence in support of an Mfd-UvrA complex in cells.

## Discussion

UvrA is the central player in NER since it performs critical functions in both GGR and TCR: First, it recognizes DNA damage as UvrA_2_ or UvrA_2_B_(2)_ and loads UvrB in GGR and second, it stimulates TCR by interacting with Mfd.

In this work, we set out to visualize the binding behaviour of fluorescently tagged UvrA in living cells over the experimentally accessible, and biologically relevant timescale of 0.1 – 1000 s. We found that in live cells, UvrA exhibits foci with residence times ranging from a few hundred milliseconds, to tens of seconds. Here, using single-molecule imaging in combination with chemical and genetic tools, we characterized the DNA-binding lifetimes of UvrA in its interactions with its key binding partners in live cells.

We identified that UvrA is long-lived on DNA in the absence of Mfd and UvrB with a lifetime of 24 s. Consistent with previous studies demonstrating that UvrB promotes the dissociation of UvrA *in vitro*, UvrA also dissociated faster in *uvrB*^+^ cells ^42, 43, 44^. The insensitivity to rif or UV treatment suggests that the short-lived population of DNA-bound UvrA with a lifetime of ∼2 s corresponds to UvrA engaged in damage search. Additionally, we also detected a long-lived species of UvrA in untreated cells. Using TCR-deficient strains and rif treatment, we assigned the lifetime of UvrA in interactions with UvrB to be 8.7 s. This lifetime likely corresponds to UvrA dissociation following loading of UvrB on DNA since the loading-deficient UvrB mutant arrests UvrA for 148 s (see Fig. 3). Overall, our findings are consistent with a previous study that detected a short-lived UvrA species in untreated cells, and a long-lived species upon UV treatment ^23^.

Three observations provide evidence for an Mfd-UvrA interaction in cells: first, cells lacking UvrB exhibited UvrA-YPet foci that were five-fold longer lived compared to cells lacking both Mfd and UvrB, suggesting that UvrA forms a highly stable complex with Mfd. Second, the long-lived lifetime of UvrA in non-UV treated cells was sensitive to rif treatment, whereby these cells lost a slowly dissociating species of UvrA upon rif treatment, presumably engaged with Mfd. Curiously, the DNA binding kinetics of UvrA in Mfd-UvrA_2_-UvrB complexes could only be distinguished from those in UvrA_2_B_2_ under conditions where UvrB was present in limiting conditions, suggesting a role for UvrB in resolving Mfd-UvrA complexes in cells (Fig. 5f). A prediction of this hypothesis is that elevated concentrations of UvrB would lead to an enhanced turnover of Mfd in the Mfd-UvrA_2_-UvrB complex. Third, experiments on cells exposed to UV light revealed that the binding kinetics of UvrA mirrored those of Mfd suggesting that these two proteins participate in a TCR handoff complex during the SOS response. Further, UvrA is turned over faster in an Mfd-dependent manner during the SOS response compared to untreated cells (Fig. 5). The elevated copy numbers of UvrA and UvrB following UV treatment (Fig. 5b and refs ^47, 48^) may explain the faster turnover of UvrA during the SOS response. The lifetime of UvrA in TCR-deficient UV-treated cells increased to 15 s, but remained low in UV-treated wild-type cells. An explanation for this striking difference may be found in work by Crowley and Hanawalt ^46^. The differences observed in our experiments may fundamentally be attributable to the efficiencies with which UvrA recognizes CPDs and 6-4 photoproducts. CPD lesions^51, 52^ being less helix distorting than 6-4 photoproducts^53, 54^, are generally inefficiently recognized by damage recognition enzymes in NER across organisms^55, 56^. CPD recognition by RNA polymerase and subsequent TCR at such sites is a major mechanism for CPD removal after UV treatment^46^. Compared to untreated cells that remove greater than 80% of CPD lesions within 40 minutes following UV treatment, rif-treated cells removed less than 50% of CPD lesions at the same time point^46^. In contrast, 6-4 photoproduct removal was uninfluenced by rif treatment^46^. Further, this report also demonstrated that the copy number of UvrAB determines whether CPDs are efficiently removed following UV treatment. Considering the low expression level of UvrA-YPet in our strain (16 ± 4 subunits per cell during normal growth), it is likely that the initial time window in which UvrA is turned over rapidly in Δ*mfd* cells reflects repair of 6-4 photoproducts. The later time windows in which UvrA exhibits a lifetime of 15 s may reflect repair of CPD lesions that are poorly detected by UvrA. Extending this argument, the prediction from the Crowley and Hanawalt study is that the rapid turnover of UvrA in wild-type cells in the 25-100 min time window reflects the repair of CPDs *via* TCR. Indeed, consistent with this prediction, we found that the binding kinetics of Mfd mirror those of UvrA in this time period. This finding is also consistent with the well-established finding that the repair of UV lesions is biphasic – a period of rapid removal of 6-4 photoproducts is followed by a slower repair of CPDs^46^.

Finally, our data suggest that the enhancement of the rate of repair in TCR *vs*. GGR measured in bulk is best explained by enhanced target search in TCR compared to GGR. This model has been previously proposed in the literature based on evidence from *in vitro* studies^6, 13^. In this model, stalled RNAP is recognized by Mfd, leading to the exposure of Mfd’s UvrB-homology module (BHM) that in turn acts as an ‘flag’ for UvrAB. Damage recognition by UvrAB would then follow initial recruitment to the site of the lesion. In contrast, target search during global genomic repair would comprise of repeated cycles of 3D diffusion of UvrA(B) to sites of undamaged DNA followed by subsequent turn-over of UvrA by UvrB until the damage surveillance complex stochastically encounters damaged DNA.

## METHODS

### Construction of strains and plasmids

*Escherichia coli* MG1655 *uvrA-YPet* was constructed using λ Red recombination as previously described for Mfd using the linker sequence described previously^12^. Deletion constructs were created by replacing the indicated gene with a kanamycin cassette flanked by FRT sites as described previously^12^. Sequence specified wild-type *uvrA* and *uvrA*(Δ131-250) geneblocks (including the native *uvrA* promoter) were ordered from IDT (Coralville, USA) and subcloned into pHH001^12^ using standard molecular biology techniques. Plasmids were sequenced on both strands prior to use. Strain expressing mutant UvrB was created using CRISPR-Cas9 assisted λ Red recombination as previously described^57, 58^ (see Supplementary Methods).

### Cell culture for imaging

Cells were imaged in quartz-top flow cells as described previously^12^. Cells were grown in 500 μL of EZ-rich defined media (Teknova, CA, US), supplemented with 0.2% (v/v) glucose in 2 mL microcentrifuge tubes at 30 °C. For experiments involving plasmid-expressed UvrA-YPet or UvrA(Δ131-250)-YPet, spectinomycin (50 μg per mL) was added to the growth media. Cells in early exponential phase were loaded in flow cells at 30 °C, followed by a constant supply of aerated EZ-rich defined media at a rate of 30 µL per min, using a syringe pump (Adelab Scientific, Australia).

### Single-molecule live-cell imaging

Single-molecule fluorescence imaging was carried out with a custom-built microscope as previously described^12^. Briefly, the microscope comprised a Nikon Eclipse Ti body, a 1.49 NA 100x objective, a 514-nm Sapphire LP laser (Coherent) operating at a power density of 71 W cm^-2^, an ET535/30m emission filter (Chroma) and a 512 x 512 pixel^2^ EM-CCD camera (either Photometrics Evolve or Andor iXon 897). The microscope was operated in near-TIRF illumination^59^ and was controlled using NIS-Elements (Nikon). PAmCherry-tagged proteins were imaged as described previously^12^.

Fluorescence images were acquired in time-series format with 100 ms frames. Each video acquisition contained two phases. The first phase (50 frames) aimed to lower background signal by continuous illuminating, causing most of the fluorophores to photo-bleach or to assume a dark state. The second phase (single-molecule phase lasting for 100 frames) is when single molecules can be reliably tracked on a low background signal. In the second phase, consecutive frames were acquired continuously or with a delay time (τ_d_).

### Image analysis

Image analysis was performed in Fiji^60^, using the Single Molecule Biophysics plugins (available at https://github.com/SingleMolecule/smb-plugins), and MATLAB. First, raw data were converted to TIF format, following by background correction and image flattening as previously described^12^. Next, foci were detected in the reactivation phase by applying a discoidal average filter (inner radius of one pixel, outer radius of three pixels), then selecting pixels above the intensity threshold. Foci detected within 3-pixel radius (318 nm) in consecutive frames were considered to belong to the same binding event.

### Interval imaging for dissociation kinetics measurements

Interval imaging was performed as described previously^12^. Briefly, the photobleaching phase contained 50 continuous 0.1-s frames. In phase II, 100 discontinuous 0.1-s frames (τ_int_ = 0.1 s) were collected with time-lapse time (τ_tl_) ranging from τ_tl_ = (0.1, 0.2, 0.5, 1, 2, 4, 8, 10). In each experiment, videos with varying τ_d_ were acquired. Foci were detected using a relative intensity threshold of 7 or 8 above the background as appropriate. Depending on the construct being imaged, between 3-15 repeats of each experiment were collected for each strain. Cumulative residence time distribution of binding events detected in all data sets were generated for each τ_tl_. Cumulative residence time distributions were globally fitting to single- or double-exponential models to obtain binding lifetimes and amplitudes of kinetic subpopulations (custom-written MATLAB codes, see ref. ^24^). Here, the choice of fitting model was decided based on the shape of *k*_eff_τ_tl_ vs. τ_tl_ plot (ref.^32^), in which the effective decay rate *k*_eff_ represents the sum of the normalized photobleaching rate *k*_b_ and the off rate *k*_off_ (*k*_eff_ = *k*_b_τ_int_ /τ_tl_ + *k*_off_) (ref. 32). The linearity of *k*_eff_τ_tl_ vs. τ_tl_ plot represents a single kinetic population whereas deviations from linear fits indicate multiple kinetic subpopulations (ref. 32). The corresponding *k*_eff_τ_tl_ vs. τ_tl_ plot was obtained as described previously^12, 24^, with the shaded error bar representing standard deviations of ten bootstrapped samples deriving from 80% of the compiled binding events (custom-written MATLAB codes)^24^. Given that this method is based on fitting mixtures of exponential models to the data, this approach reliably resolves off rates that are at least three-fold apart. In experiments involving rifampicin treatment, cells were incubated in growth media containing rifampicin (50 μg per mL) for 30 min in the flow cell prior to imaging.

### Experiments involving UV irradiation

UV survival assays were performed as described previously in references^12, 61^. UV irradiation was delivered *in situ* as described previously^61^. The UV flux was measured prior to UV irradiation, and the exposure time was adjusted to provide a dose of 20 Jm^-2^. For experiments involving interval imaging following UV exposure, τ_tl_ = (0.1, 0.3, 1, 3, 10) values were used to minimize the time taken to complete one round of interval imaging.

## ACKNOWLEDGEMENTS

H.G. acknowledges the University of Wollongong and the Faculty of Science, Medicine and Health for funding of the Andor iXon 897 used in this work. A.M. van Oijen would like to acknowledge support by the Australian Research Council (DP150100956 and FL140100027).

## AUTHOR CONTRIBUTIONS

Construct creation: H.G. and H.N.H.; Data curation: H.G.; Data analysis: H.G. and H.N.H.; Software: H.N.H. and H.G.; Writing—Original draft: H.G.; Writing—review and editing: H.N.H., H.G. and A.M.v.O.; Conceptualization: H.G.; Supervision: H.G. and A.M.v.O.

### Data availability

The data that support the findings of this study are available from the corresponding author upon reasonable request.

### Code availability

Custom code used for data analyses has been made publicly available at github.com/hghodke/bacterial_live_cell_interval_imaging

