## Supplementary Information for "Single-molecule live-cell imaging visualizes parallel pathways of prokaryotic nucleotide excision repair"

by

Harshad Ghodke, Han N. Ho, Antoine M. van Oijen

**This PDF file includes:**

Supplementary Figures 1 to 8
Supplementary Tables 1 to 4
Supplementary Notes 1 to 3
Supplementary References

### Supplementary Figures

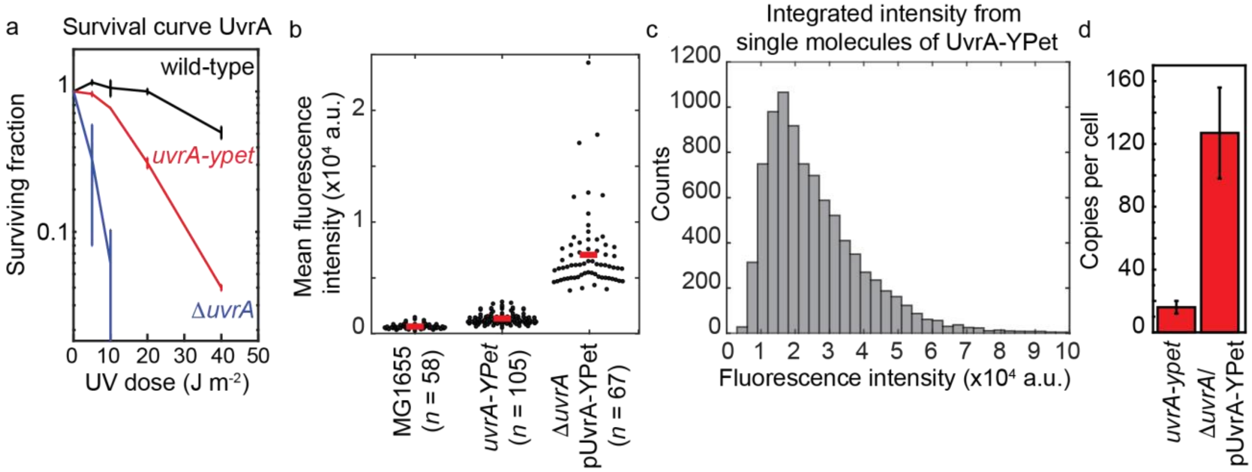


**Supplementary Figure 1:**

a. Survival assay showing survival of wild-type (black), *uvrA-YPet* (red) and Δ*uvrA* (blue) cells upon exposure to 20 Jm$^{-2}$ of 254-nm UV light. Error bars represent standard error of the mean from two independent experiments, each experiment involved three technical replicates.

b. Mean fluorescence intensity of wild-type MG1655, *uvrA-YPet* cells and cells carrying a low-copy plasmid expressing UvrA-YPet upon excitation with 514-nm light. Red line indicates mean of the distribution. *n*, number of cells.

c. Histogram of integrated fluorescence intensities of single UvrA-YPet foci in *uvrA-YPet* cells upon excitation with 514-nm light.

d. Bar plots show mean copy number of UvrA-YPet measured in 105 *uvrA-YPet* cells and 67 Δ*uvrA*/pUvrA-YPet cells. Error bars are standard deviations.

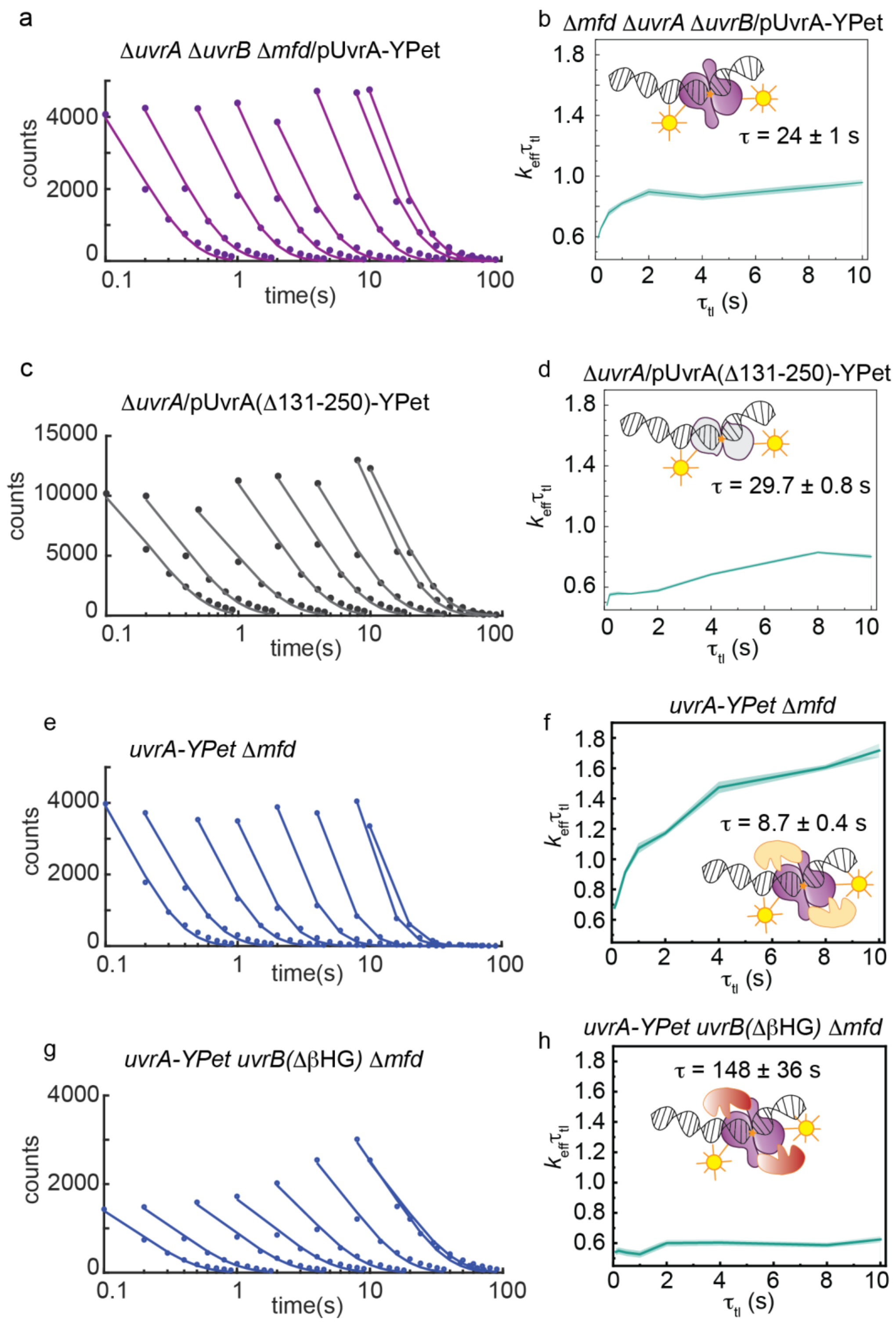
a
ΔuvrA ΔuvrB Δmfd/pUvrA-YPet
counts
time(s)
b
Δmfd ΔuvrA ΔuvrB/pUvrA-YPet
τ = 24 ± 1 s
$k_{eff}\tau_{tl}$
$\tau_{tl}$ (s)
c
ΔuvrA/pUvrA(Δ131-250)-YPet
counts
time(s)
d
ΔuvrA/pUvrA(Δ131-250)-YPet
τ = 29.7 ± 0.8 s
$k_{eff}\tau_{tl}$
$\tau_{tl}$ (s)
e
uvrA-YPet Δmfd
counts
time(s)
f
uvrA-YPet Δmfd
τ = 8.7 ± 0.4 s
$k_{eff}\tau_{tl}$
$\tau_{tl}$ (s)
g
uvrA-YPet uvrB(ΔβHG) Δmfd
counts
time(s)
h
uvrA-YPet uvrB(ΔβHG) Δmfd
τ = 148 ± 36 s
$k_{eff}\tau_{tl}$
$\tau_{tl}$ (s)

**Supplementary Figure 2:**
a. Cumulative residence time distributions (CRTDs, circles) obtained from interval imaging of UvrA-YPet in Δ*uvrA* Δ*uvrB* Δ*mfd* cells. Lines are mono-exponential fits to CRTDs.
b. The $k_{eff}\tau_{tl}$ plot obtained from fitting CRTDs of UvrA-YPet in Δ*uvrA* Δ*uvrB* Δ*mfd* cells. Shaded error bands are standard deviations from ten bootstrapped samples. Cartoon (inset) illustrates UvrA-YPet (purple) in complex with DNA.
c. CRTDs (circles) obtained from interval imaging of UvrA(Δ131-250)-YPet in Δ*uvrA* cells. Lines are mono-exponential fits to CRTDs.
d. The $k_{eff}\tau_{tl}$ plot obtained from fitting CRTDs of UvrA(Δ131-250)-YPet in Δ*uvrA* cells. Shaded error bands are standard deviations from ten bootstrapped samples. Cartoon (inset) illustrates c UvrA(Δ131-250)-YPet (grey) in complex with DNA.
e. CRTDs (circles) obtained from interval imaging of UvrA-YPet in *uvrA-YPet* Δ*mfd* cells. Lines are mono-exponential fits to CRTDs.
f. The $k_{eff}\tau_{tl}$ plot obtained from fitting CRTDs of UvrA-YPet in *uvrA-YPet* Δ*mfd* cells. Shaded error bands are standard deviations from ten bootstrapped samples. Cartoon (inset) illustrates the complex formed by UvrA-YPet (purple) and UvrB (orange) with DNA.
g. CRTDs (circles) obtained from interval imaging of UvrA-YPet in *uvrA-YPet uvrB(*Δβ*HG)* Δ*mfd* cells. Lines are mono-exponential fits to CRTDs.
h. The $k_{eff}\tau_{tl}$ plot obtained from fitting CRTDs of UvrA-YPet in *uvrA-YPet uvrB(*Δβ*HG)* Δ*mfd* cells. Shaded error bands are standard deviations from ten bootstrapped samples. Cartoon (inset) illustrates the complex formed by UvrA-YPet (purple) and UvrB(ΔβHG) (red) with DNA.

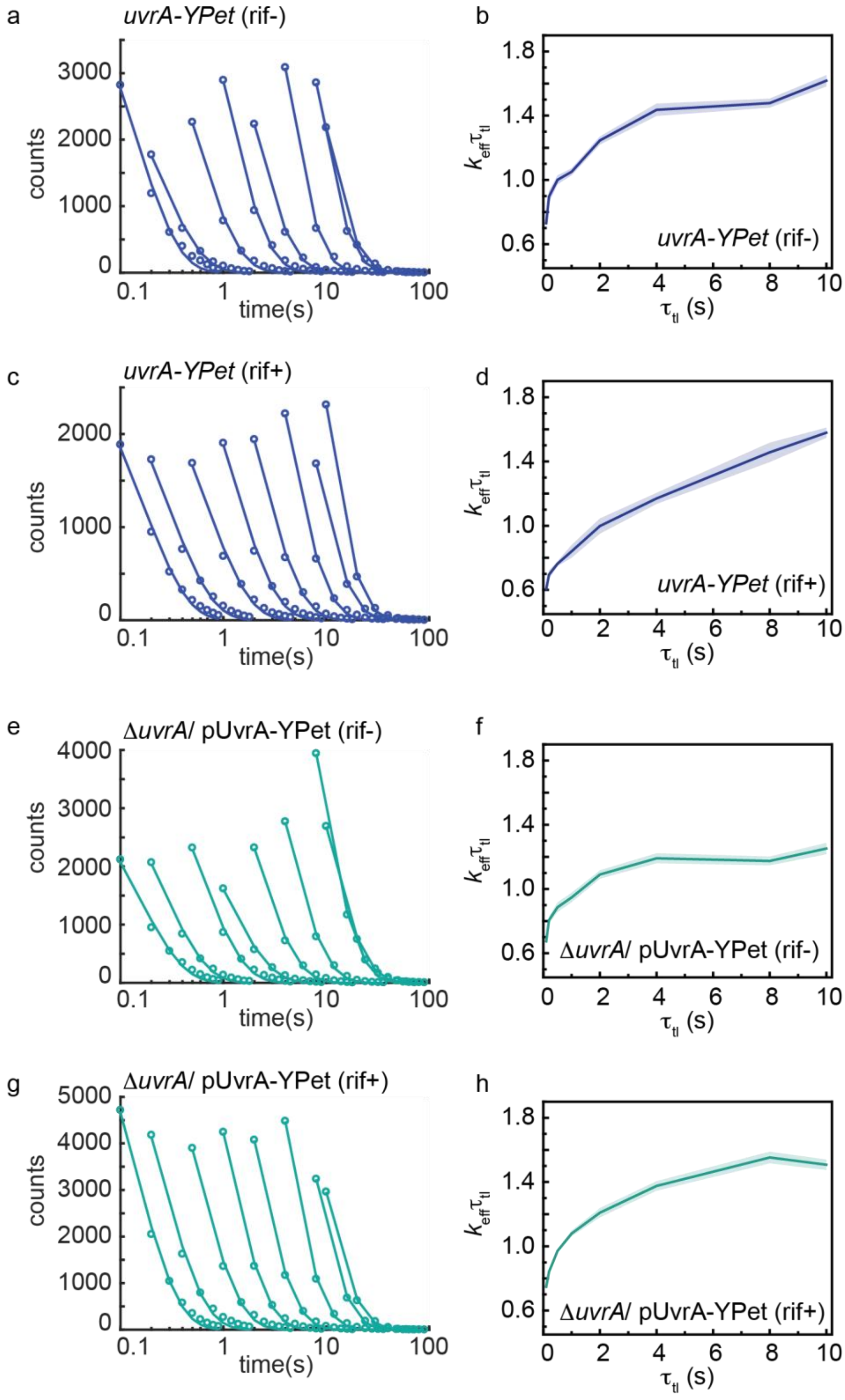

a
uvrA-YPet (rif-)
counts
time(s)
b
$k_{eff}\tau_{tl}$
uvrA-YPet (rif-)
$\tau_{tl}$ (s)
c
uvrA-YPet (rif+)
counts
time(s)
d
$k_{eff}\tau_{tl}$
uvrA-YPet (rif+)
$\tau_{tl}$ (s)
e
ΔuvrA/ pUvrA-YPet (rif-)
counts
time(s)
f
$k_{eff}\tau_{tl}$
ΔuvrA/ pUvrA-YPet (rif-)
$\tau_{tl}$ (s)
g
ΔuvrA/ pUvrA-YPet (rif+)
counts
time(s)
h
$k_{eff}\tau_{tl}$
ΔuvrA/ pUvrA-YPet (rif+)
$\tau_{tl}$ (s)

**Supplementary Figure 3:**
a. CRTDs (circles) obtained from interval imaging of UvrA-YPet in *uvrA-YPet* cells. Lines are mono-exponential fits to CRTDs.
b. The $k_{eff}\tau_{tl}$ plot obtained from fitting CRTDs of UvrA-YPet in *uvrA-YPet* cells. Shaded error bands are standard deviations from ten bootstrapped samples.
c. CRTDs (circles) obtained from interval imaging of UvrA-YPet in rif-treated *uvrA-YPet* cells. Lines are mono-exponential fits to CRTDs.
d. The $k_{eff}\tau_{tl}$ plot obtained from fitting CRTDs of UvrA-YPet in rif-treated *uvrA-YPet* cells. Shaded error bands are standard deviations from ten bootstrapped samples.
e. CRTDs (circles) obtained from interval imaging of UvrA-YPet in Δ*uvrA*/pUvrA-YPet cells. Lines are mono-exponential fits to CRTDs.
f. The $k_{eff}\tau_{tl}$ plot obtained from fitting CRTDs of UvrA-YPet in Δ*uvrA*/pUvrA-YPet cells. Shaded error bands are standard deviations from ten bootstrapped samples.
g. CRTDs (circles) obtained from interval imaging of UvrA-YPet in rif-treated Δ*uvrA*/pUvrA-YPet cells. Lines are mono-exponential fits to CRTDs.
h. The $k_{eff}\tau_{tl}$ plot obtained from fitting CRTDs of UvrA-YPet in rif-treated Δ*uvrA*/pUvrA-YPet cells. Shaded error bands are standard deviations from ten bootstrapped samples.

*uvrA-YPet ∆mfd* (UV 20 $Jm^{-2}$)

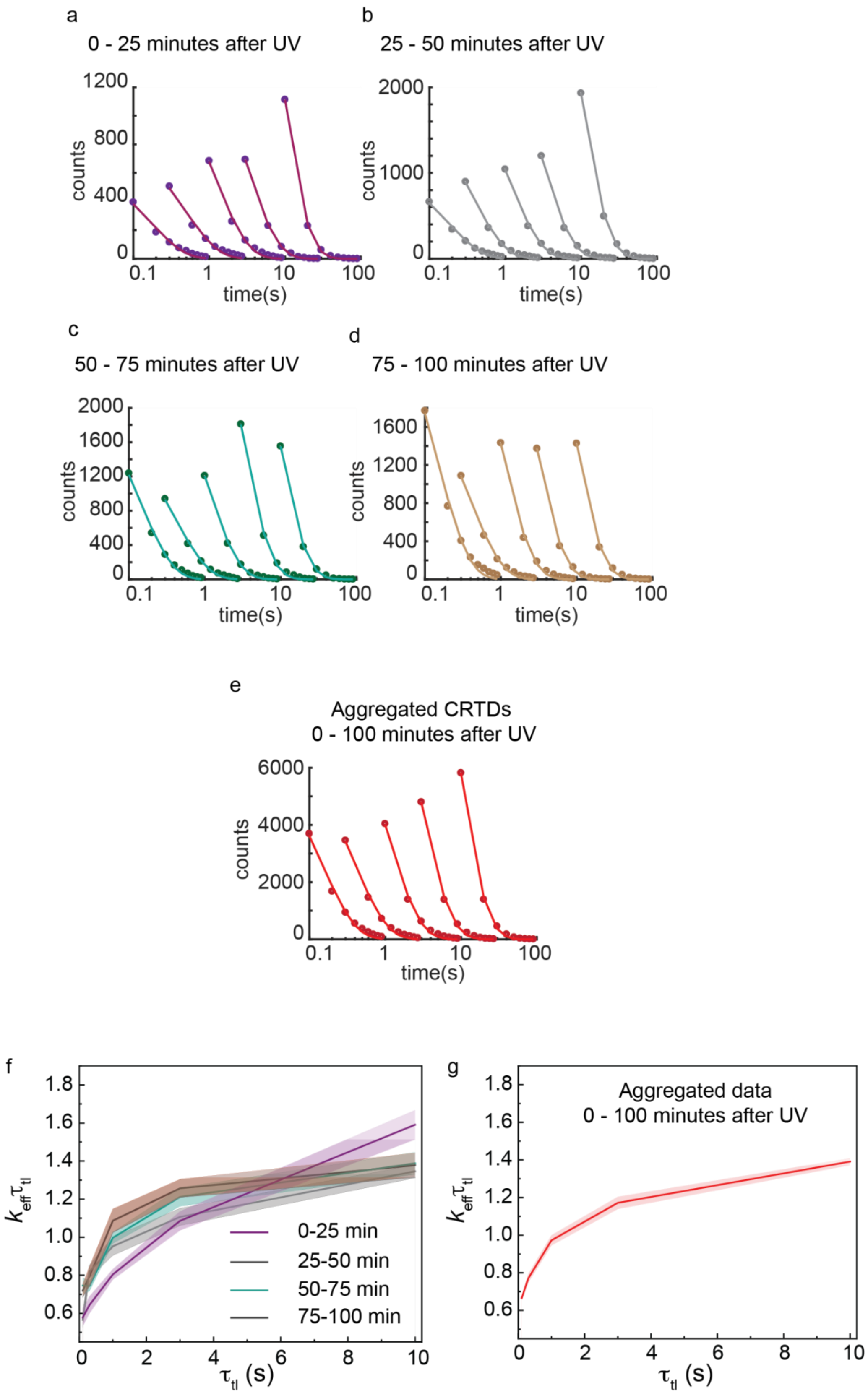

**Supplementary Figure 4:**

a. CRTDs (circles) obtained from interval imaging of UvrA-YPet in *uvrA-YPet* Δ*mfd* cells within the first 25 minutes following exposure to a 20 $Jm^{-2}$ dose of 254-nm light. Lines are mono-exponential fits to CRTDs.

b. CRTDs (circles) obtained from interval imaging of UvrA-YPet in *uvrA-YPet* Δ*mfd* cells 25-50 minutes following exposure to a 20 $Jm^{-2}$ dose of 254-nm light. Lines are mono-exponential fits to CRTDs.

c. CRTDs (circles) obtained from interval imaging of UvrA-YPet in *uvrA-YPet* Δ*mfd* cells 50-75 minutes following exposure to a 20 $Jm^{-2}$ dose of 254-nm light. Lines are mono-exponential fits to CRTDs.

d. CRTDs (circles) obtained from interval imaging of UvrA-YPet in *uvrA-YPet* Δ*mfd* cells 75-100 minutes following exposure to a 20 $Jm^{-2}$ dose of 254-nm light. Lines are mono-exponential fits to CRTDs.

e. CRTDs (circles) obtained from interval imaging of UvrA-YPet in *uvrA-YPet* Δ*mfd* cells within the first 100 minutes following exposure to a 20 $Jm^{-2}$ dose of 254-nm light. Lines are mono-exponential fits to CRTDs.

f. The $k_{eff}\tau_{tl}$ plots obtained from fitting CRTDs of UvrA-YPet in UV-treated *uvrA-YPet* Δ*mfd* cells as a function of time following UV exposure. Shaded error bands are standard deviations from ten bootstrapped samples. Purple, 0-25 minutes; grey, 25-50 minutes; cyan, 50-75 minutes; brown, 75-100 minutes.

g. The $k_{eff}\tau_{tl}$ plot obtained from fitting CRTDs of UvrA-YPet in UV-treated *uvrA-YPet* Δ*mfd* cells within the first 100 minutes following UV exposure.

***uvrA-YPet* (UV 20 Jm$^{-2}$)**

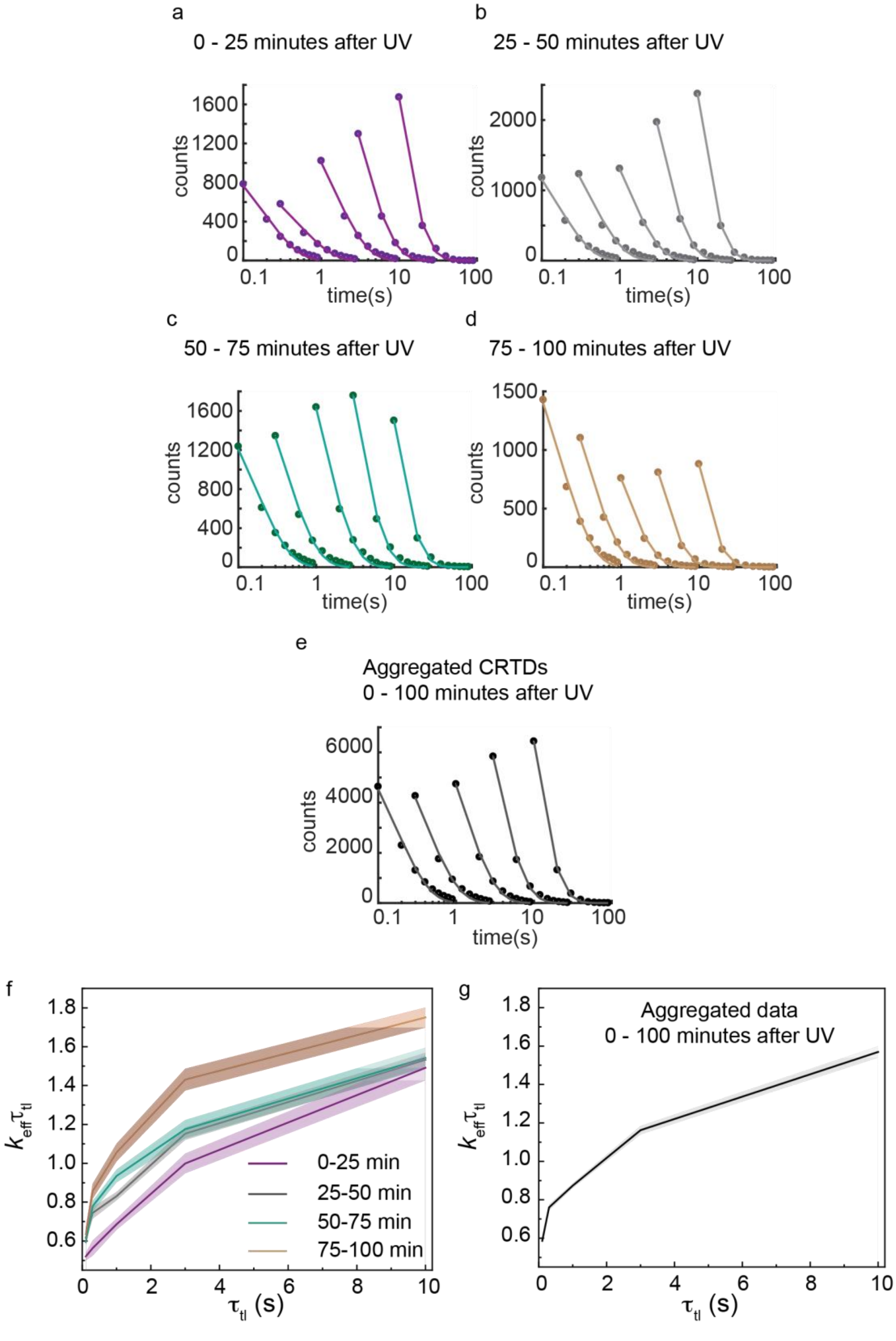

**Supplementary Figure 5:**

a. CRTDs (circles) obtained from interval imaging of UvrA-YPet in *uvrA-YPet* cells within the first 25 minutes following exposure to a 20 $Jm^{-2}$ dose of 254-nm light. Lines are mono-exponential fits to CRTDs.

b. CRTDs (circles) obtained from interval imaging of UvrA-YPet in *uvrA-YPet* cells 25-50 minutes following exposure to a 20 $Jm^{-2}$ dose of 254-nm light. Lines are mono-exponential fits to CRTDs.

c. CRTDs (circles) obtained from interval imaging of UvrA-YPet in *uvrA-YPet* cells 50-75 minutes following exposure to a 20 $Jm^{-2}$ dose of 254-nm light. Lines are mono-exponential fits to CRTDs.

d. CRTDs (circles) obtained from interval imaging of UvrA-YPet in *uvrA-YPet* cells 75-100 minutes following exposure to a 20 $Jm^{-2}$ dose of 254-nm light. Lines are mono-exponential fits to CRTDs.

e. CRTDs (circles) obtained from interval imaging of UvrA-YPet in *uvrA-YPet* cells within the first 100 minutes following exposure to a 20 $Jm^{-2}$ dose of 254-nm light. Lines are mono-exponential fits to CRTDs.

f. The $k_{\text{eff}}\tau_{\text{tl}}$ plots obtained from fitting CRTDs of UvrA-YPet in UV-treated *uvrA-YPet* cells as a function of time following UV exposure. Shaded error bands are standard deviations from ten bootstrapped samples. Purple, 0-25 minutes; grey, 25-50 minutes; cyan, 50-75 minutes; brown, 75-100 minutes.

g. The $k_{\text{eff}}\tau_{\text{tl}}$ plot obtained from fitting CRTDs of UvrA-YPet in UV-treated *uvrA-YPet* cells within the first 100 minutes following UV exposure.

***mfd-YPet* (UV 20 Jm$^{-2}$)**

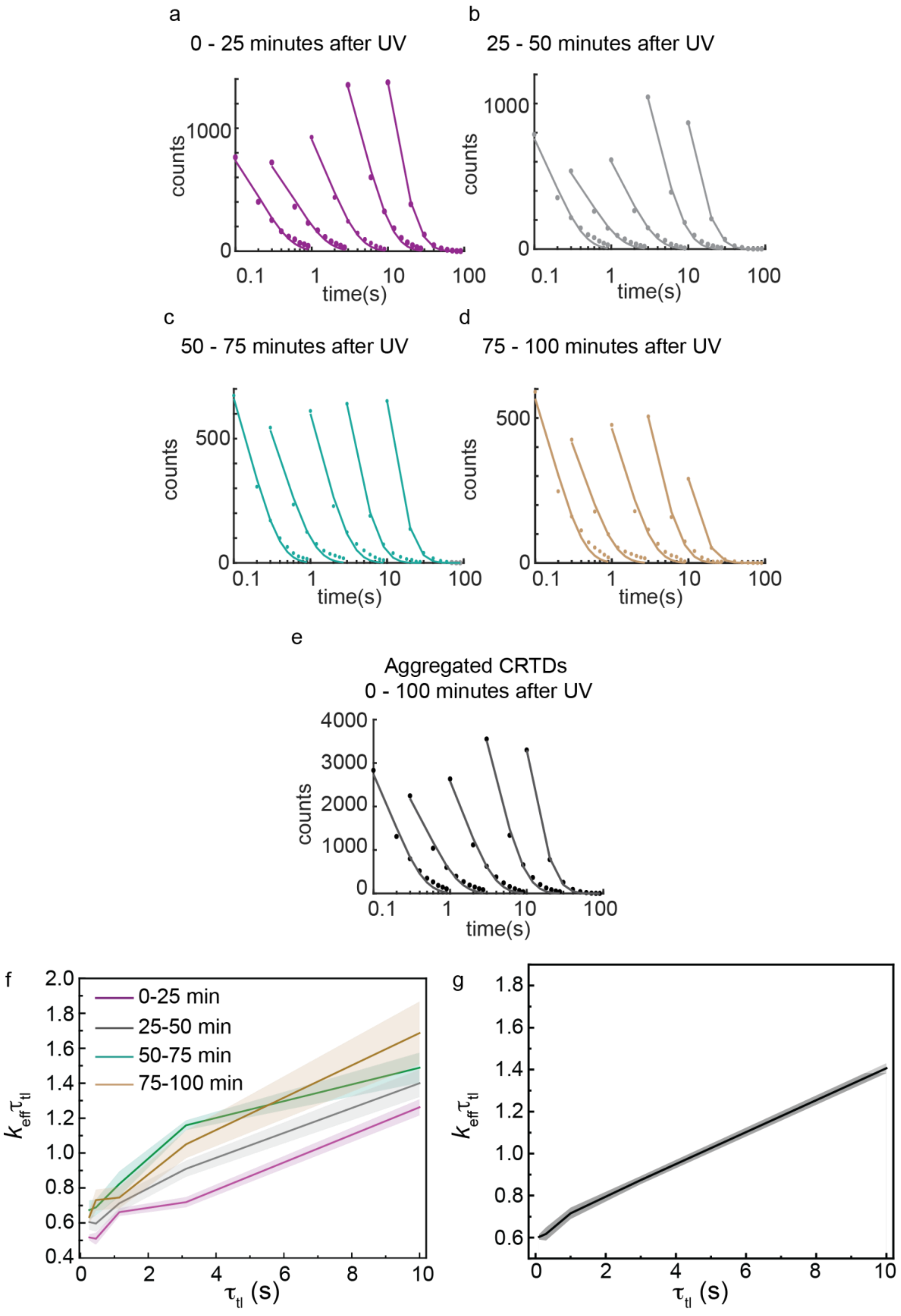

**Supplementary Figure 6:**

a. CRTDs (circles) obtained from interval imaging of Mfd-YPet in *mfd-YPet* cells within the first 25 minutes following exposure to a 20 $Jm^{-2}$ dose of 254-nm light. Lines are mono-exponential fits to CRTDs.

b. CRTDs (circles) obtained from interval imaging of Mfd-YPet in *mfd-YPet* cells 25-50 minutes following exposure to a 20 $Jm^{-2}$ dose of 254-nm light. Lines are mono-exponential fits to CRTDs.

c. CRTDs (circles) obtained from interval imaging of Mfd-YPet in *mfd-YPet* cells 50-75 minutes following exposure to a 20 $Jm^{-2}$ dose of 254-nm light. Lines are mono-exponential fits to CRTDs.

d. CRTDs (circles) obtained from interval imaging of Mfd-YPet in *mfd-YPet* cells 75-100 minutes following exposure to a 20 $Jm^{-2}$ dose of 254-nm light. Lines are mono-exponential fits to CRTDs.

e. CRTDs (circles) obtained from interval imaging of Mfd-YPet in *mfd-YPet* cells within the first 100 minutes following exposure to a 20 $Jm^{-2}$ dose of 254-nm light. Lines are mono-exponential fits to CRTDs.

f. The $k_{eff}\tau_{tl}$ plots obtained from fitting CRTDs of Mfd-YPet in UV-treated *mfd-YPet* cells as a function of time following UV exposure. Shaded error bands are standard deviations from ten bootstrapped samples. Purple, 0-25 minutes; grey, 25-50 minutes; cyan, 50-75 minutes; brown, 75-100 minutes.

g. The $k_{eff}\tau_{tl}$ plot obtained from fitting CRTDs of Mfd-YPet in UV-treated *mfd-YPet* cells within the first 100 minutes following UV exposure.

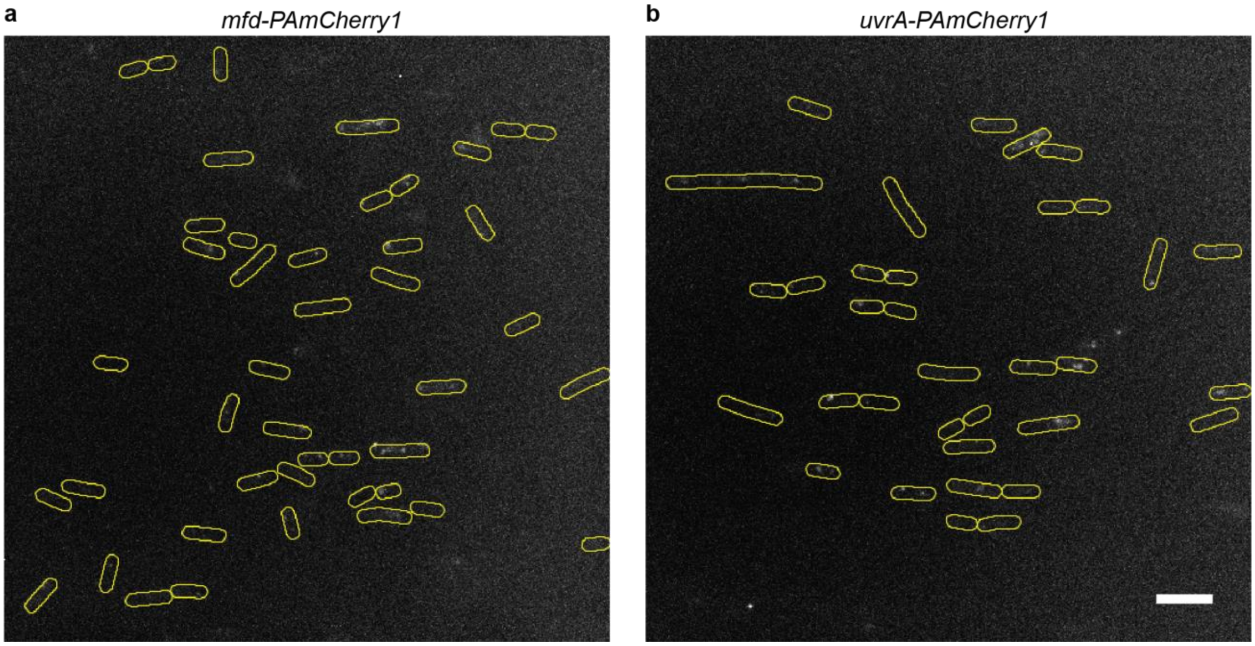


**Supplementary Figure 7:**
a. Maximum intensity projection of *mfd-PAmCherry1* cells obtained upon exposure to 405-nm and 568-nm light.
b. Maximum intensity projection of *uvrA-PAmCherry1* cells obtained upon exposure to 405-nm and 568-nm light.
Note the significantly fewer localizations obtained in these strains compared to the YPet tagged constructs (Figure 2) and reference 1. Cell outlines (yellow) are provided as guide to the eye. Scale bar represents 5 μm.

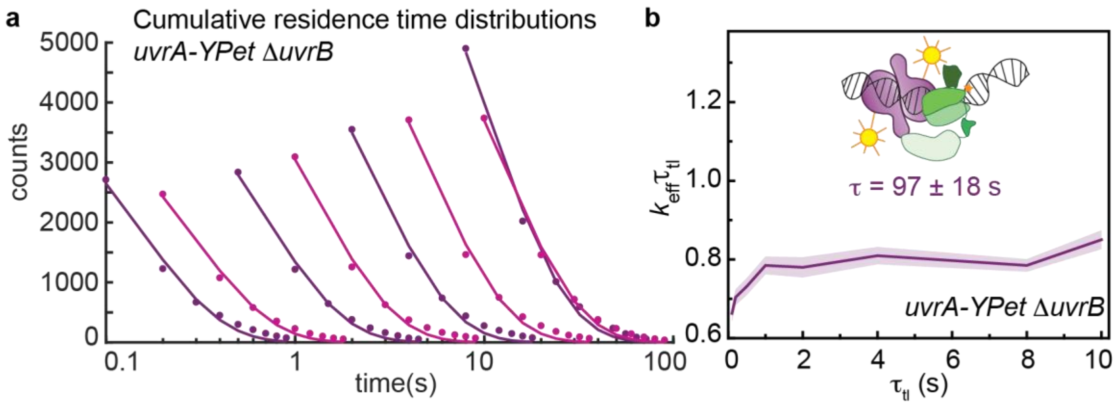


**Supplementary Figure 8:**
a. CRTDs (circles) obtained from interval imaging of UvrA-YPet in *uvrA-YPet* Δ*uvrB* cells. Lines are mono-exponential fits to CRTDs.
b. The $k_{eff}\tau_{tl}$ plot obtained from fitting CRTDs of UvrA-YPet in *uvrA-YPet* Δ*uvrB* cells. Shaded error bands are standard deviations from ten bootstrapped samples. Cartoon (inset) illustrates the arrested complex formed by UvrA-YPet (purple) and Mfd (green) in *uvrA-YPet* Δ*uvrB* cells.

### Supplementary Tables

**Supplementary Table 1.** Bacterial strains. All strains are in *E. coli* K-12 MG1655 background.

| **Strain/genotypes** | **Source/Technique** |
|---|---|
| *uvrA-YPet* | This study/ λ Red recombination |
| *uvrA-PAmCherry1* | This study/ λ Red recombination |
| *uvrA::kanR* | This laboratory |
| *uvrB::kanR* | This study/ λ Red recombination |
| *mfd::kanR* | This laboratory |
| *ΔuvrA Δmfd ΔuvrB* | This study |
| *ΔuvrA/*pUvrA-YPet | This study |
| *ΔuvrA Δmfd ΔuvrB/*pUvrA-YPet | This study |
| *ΔuvrA Δmfd ΔuvrB/* pUvrA(Δ131-250)-YPet | This study |
| *uvrA-YPet Δmfd* | This study/ P1 transduction |
| *uvrA-YPet uvrB(ΔβHG)* | This study/ CRISPR-Cas9 assisted λ Red recombination |
| *uvrA-YPet uvrB(ΔβHG) Δmfd* | This study/ P1 transduction |
| *uvrA-YPet ΔuvrB* | This study/ P1 transduction |
| *mfd-YPet* | This laboratory |
| *mfd-PAmCherry1* | This study |

**Supplementary Table 2.** Oligonucleotides used for colony PCR and CRISPR-Cas9 assisted λ Red recombination.

| Oligo names | Sequence |
|---|---|
| *Colony PCR for Cas9 verification* | |
| dCas9dL5_303_F | CAGACCGCCACAGTATCAAA |
| pCas9_6700_R | GGAAGGTATCCGACTGCTG |
| *Cloning of pCRISPR variants* | |
| pCRISPR_UvrB_Y96A_S | AAA CCC TAC TAC GAC TAC TAT CAG CG |
| pCRISPR_UvrB_Y96A_AS | AAA ACG CTG ATA GTA GTC GTA GTA GG |
| *Recombineric ssDNA* | |
| UvrB_ΔβHG_ ssDNA | CAA TAT GTT CGT TAA CCG AGG CAT CTT TCT CAA TGA AAG TGC CAT AAT AGT CGT AGT AGG AAA CGA AAT ATT CCA CCG CGT TTT CCG |

**Supplementary Table 3.** Plasmids.

| Plasmid | Source/Technique |
|---|---|
| pHH001 (pSC101-based) | This laboratory[1] |
| pUvrA-YPet | This study/Sub-cloning into pHH001 |
| pUvrA(Δ131-250)-YPet | This study/Sub-cloning into pHH001 |
| pKD46 | Cox lab[3] |
| pCas9 | Addgene # 42876, Marraffini lab[4] |
| pCRISPR | Addgene # 42875, Marraffini lab[4] |
| pCRISPR-UvrB-Y96A | This study/Sub-cloning into pCRISPR |

**Supplementary Table 4.** Global fitting outputs for measurements of residence time of UvrA-YPet and mutants in various genetic backgrounds. Bootstrapped CRTDs were fitted two single- and bi-exponential models (model 1 and 2 respectively). Selection of model was carried out as in ref.[1]. The chosen model and fitting outcomes are highlighted. $k_b$, the photobleaching rate, $\tau_1$, the slow lifetime, $B$, the amplitude of the slowly dissociating population, $\tau_2$, the fast lifetime.

| Genetic background | Model | $k_b$ ± Error ($s^{-1}$) | $\tau_1$ ± Error (s) | $B$ ± Error (%) | $\tau_2$ ± Error (s) |
|---|---|---|---|---|---|
| Δ*uvrA* Δ*uvrB* Δ*mfd*/ pUvrA-YPet | 1 | 7.0 ± 0.1 | 33 ± 2 | 100 | - |
| | **2** | **5.5 ± 0.1** | **24 ± 1** | **28 ± 2** | **1.6 ± 0.2** |
| Δ*uvrA*/ pUvrA(Δ131-250)-YPet | **1** | **5.2 ± 0.1** | **29.7 ± 0.8** | **100** | **-** |
| | 2 | 4.3 ± 0.1 | 21.9 ± 0.5 | 47 ± 1 | 0.75 ± 0.06 |
| *uvrA-YPet* Δ*mfd* | 1 | 7.7 ± 0.1 | 7.4 ± 0.2 | 100 | - |
| | **2** | **6.3 ± 0.2** | **8.7 ± 0.4** | **22 ± 2** | **1.5 ± 0.1** |
| *uvrA-YPet uvrB(ΔBHG)* Δ*mfd* | **1** | **5.5 ± 0.1** | **148 ± 36** | **100** | **-** |
| | 2 | 4.7 ± 0.1 | 852 ± 272 | 42 ± 3 | 5.7 ± 0.6 |
| *uvrA-YPet* Δ*mfd* 0-100 min UV+ | 1 | 7.9 ± 0.2 | 14.0 ± 0.7 | 100 | - |
| | **2** | **6.2 ± 0.2** | **13.1 ± 0.6** | **23 ± 3** | **1.6 ± 0.2** |
| *uvrA-YPet* Δ*mfd* 0-25 min UV+ | 1 | 6.8 ± 0.2 | 10.8 ± 0.8 | 100 | - |
| | **2** | **5.0 ± 0.6** | **10.1 ± 1.5** | **27 ± 7** | **1.5 ± 0.3** |
| *uvrA-YPet* Δ*mfd* 25-50 min UV+ | 1 | 8.1 ± 0.2 | 17.6 ± 1.1 | 100 | - |
| | **2** | **5.8 ± 0.4** | **13.5 ± 0.7** | **21 ± 4** | **1.5 ± 0.2** |
| *uvrA-YPet* Δ*mfd* 50-75 min UV+ | 1 | 8.1 ± 0.2 | 13.3 ± 0.9 | 100 | - |
| | **2** | **6.5 ± 0.2** | **15 ± 2** | **24 ± 3** | **1.8 ± 0.3** |
| *uvrA-YPet* Δ*mfd* 75-100 min UV+ | 1 | 8.0 ± 0.2 | 11 ± 1 | 100 | - |
| | **2** | **6.5 ± 0.5** | **15 ± 4** | **17 ± 4** | **1.5 ± 0.3** |
| *uvrA-YPet* | 1 | 8.7 ± 0.2 | 10.0 ± 0.6 | 100 | - |
| | **2** | **7.2 ± 0.2** | **12.0 ± 0.8** | **21 ± 2** | **1.9 ± 0.2** |
| *uvrA-YPet* Rif-treated | 1 | 6.8 ± 0.2 | 9.4 ± 0.4 | 100 | - |
| | **2** | **5.7 ± 0.2** | **9.6 ± 0.6** | **37 ± 3** | **1.5 ± 0.3** |
| Δ*uvrA*/ pUvrA-YPet | 1 | 8.4 ± 0.1 | 21 ± 2 | 100 | - |
| | **2** | **6.8 ± 0.1** | **19 ± 1** | **26 ± 2** | **2.0 ± 0.1** |
| Δ*uvrA*/pUvrA-YPet Rif-treated | 1 | 8.5 ± 0.1 | 9.0 ± 0.2 | 100 | - |
| | **2** | **7.3 ± 0.1** | **11.5 ± 0.6** | **25 ± 2** | **1.7 ± 0.1** |
| *uvrA-YPet* 0-100 min UV+ | 1 | 6.9 ± 0.1 | 9.2 ± 0.4 | 100 | - |
| | **2** | **5.6 ± 0.1** | **10.0 ± 0.4** | **26 ± 2** | **1.6 ± 0.1** |
| *uvrA-YPet* 0-25 min UV+ | **1** | **5.7 ± 0.2** | **9.6 ± 0.8** | **100** | **-** |
| | 2 | 4.8 ± 0.1 | 12.7 ± 2.6 | 29 ± 14 | 2.6 ± 0.7 |
| *uvrA-YPet* 25-50 min UV+ | **1** | **7.2 ± 0.2** | **10.7 ± 0.5** | **100** | **-** |
| | 2 | 6.0 ± 0.3 | 11.6 ± 0.9 | 30 ± 3 | 2.2 ± 0.4 |
| *uvrA-YPet* 50-75 min UV+ | **1** | **7.2 ± 0.2** | **8.8 ± 1.0** | **100** | **-** |
| | 2 | 5.6 ± 0.2 | 10.2 ± 0.6 | 22 ± 3 | 1.4 ± 0.1 |
| *uvrA-YPet* 75-90 min UV+ | 1 | 6.9 ± 0.2 | 5.1 ± 0.5 | 100 | - |
| | **2** | **5.8 ± 0.4** | **8.4 ± 0.6** | **13 ± 3** | **1.2 ± 0.2** |

### Supplementary Notes

#### Supplementary Note 1

Sequence verified recombinant strains carrying Mfd-PAmCherry1 and UvrA-PAmCherry1 were then screened by imaging at the single-cell level. Unlike their YPet tagged counterparts, the PAmCherry1 labelled proteins were poorly expressed in MG1655 (Supplementary Fig. 7).

**Supplementary Note 2: Strain construction**

The chromosomal fusions of *uvrA-YPet* and *uvrA-PAmCherry1* and *mfd-PAmCherry1* were created using λ Red recombination as previously described[3]. Briefly, gene fragments encoding YPet or PAmCherry1, and a kanamycin resistance cassette were transformed into MG1655 cells expressing the λ Red proteins from pKD46. These gene fragments are flanked by about 40-bp sequence homologies to those immediately downstream of *uvrA* or *mfd* allele.

Knock-out strains of *uvrA::KanR*, *uvrB::KanR* and *mfd::KanR* were created *de novo*, using λ Red recombination as mentioned above. The kanamycin cassette was subsequently removed to facilitate the production of the triple-deletion strain Δ*uvrA* Δ*uvrB* Δ*mfd* via P1 transduction.

Derivatives of *uvrA-YPet* expressing mutant UvrB(ΔβHG) from the chromosome were constructed using scar-less CRISPR-Cas9 assisted λ Red recombination as previously described[4,5]. Briefly, *uvrA-YPet* electrocompetent cells were transformed with pKD46[3] and pCas9[4]. The transformants were plated on LB plate containing ampicillin (50 µg per mL) and chloramphenicol (25 µg per mL) and were incubated at 30 °C overnight. As pCas9 is prone to recombination events[4], the resulting colonies were screened with colony PCR to confirm the presence of the full-length Cas9 gene, using primers (Supplementary Table 2) targeting upstream and downstream of the Cas9 gene in pCas9.

Next, *uvrA-YPet* cells harbouring pKD46 and pCas9 were made electrocompetent. First, the cells were grown at 30 °C with shaking at 200 rpm in 50 mL of LB containing ampicillin (50 µg per mL) and chloramphenicol (25 µg per mL). In these cells, Cas9 was constitutively expressed. The expression of λ Red recombination proteins was induced with 0.2% L-arabinose (w/v) when cells reached a density around 0.4 ($OD_{600}$). When the optical density reached 0.8, cells were pelleted at 4 °C and washed twice with ice-cold water, with an additional wash using 10% glycerol. Finally, aliquots containing 40 µL of cells in microcentrifuge tubes were snap-freezed in liquid nitrogen and stored at -80 °C.

Cas9 endonuclease activities were targeted to the vicinity of the desired point mutations on the *E. coli* chromosome with the help of the guide RNAs. These guide RNAs were designed such that each contains a 20-nt complementary sequence to that on the *E. coli* chromosome 5′ of the PAM sequence (5’-NGG). They were expressed from pCRISPR-UvrB-Y96A (Supplementary Table 3), which we created following protocols from reference[4].

The point mutations were introduced by recombining the foreign single-stranded DNA (ssDNA) (Supplementary Table 2). The ssDNAs were 80- to 90-nt oligos flanked by about 40-nt of sequence homologies to the *E. coli* chromosome on both sides of the desired mutations. Base changes were selected to be within five bases from the PAM sequence. This region, termed the seed region, is the most critical to Cas9 binding and disruption in this area ensure cells harbouring the point mutations would not become target for Cas9 cutting.

Aliquots of cells expressing pCas9 and λ Red recombination proteins were transformed with 30 ng of pCRISRP-UvrB-Y96A plasmid and 500 ng of ssDNA. Positives were selected on LB plates containing 50 µg per mL of kanamycin and 25 µg per mL of chloramphenicol at 37 °C, and were screened by colony PCR, and the promoter and gene sequences were verified. Curing of pCRISPR-UvrB-Y96A was performed by propagating cells on LB plates for a week at 42 °C. This is critical for subsequent rounds of P1 transduction to create *uvrA-YPet uvrB(ΔβHG) Δmfd*.

**Supplementary Note 3: Plasmid construction**

pUvrA-YPet and pUvrA(Δ131-250)-YPet were constructed by sub-cloning the *uvrA* promoter and *uvrA* or *uvrA(Δ131-250)* allele (synthetic dsDNA, IDT, Illinois, US) into pHH001[1] at *Apa*I and *AsiS*I sites. The promoter sequence was identified as 210 nucleotides directly upstream of the *uvrA* allele on the chromosome[6].
